# Beyond Basic Characterization and Omics: Immunomodulatory Roles of Platelet-Derived Extracellular Vesicles Unveiled by Functional Testing

**DOI:** 10.1101/2023.12.31.570750

**Authors:** Mari Palviainen, Johanna Puutio, Rikke Halse Østergaard, Johannes A. Eble, Katariina Maaninka, Joseph Ndika, Otto K. Kari, Masood Kamali-Moghaddam, Kasper Kjaer-Sorensen, Claus Oxvig, Ana M Aransay, Juan Falcon-Perez, Antonio Federico, Dario Greco, Saara Laitinen, Yuya Hayashi, Pia RM Siljander

## Abstract

Renowned for their role in hemostasis and thrombosis, platelets are also increasingly recognized for their contribution in innate immunity, immunothrombosis and inflammatory diseases. Platelets express a wide range of receptors, which allows them to reach a variety of activation endpoints and grants them immunomodulatory functions. Activated platelets release extracellular vesicles (PEVs), whose formation and molecular cargo has been shown to depend on receptor-mediated activation and environmental cues.

This study compares the immunomodulatory profiles of PEVs generated via activation of platelets by different receptors, glycoprotein VI, C-type lectin-like receptor 2, and combining all thrombin-collagen receptors. Functional assays *in vivo* in zebrafish and *in vitro* in human macrophages respectively highlighted distinct homing and secretory responses triggered by the PEVs. In contrast, omics analyses of protein and miRNA cargo combined with physicochemical particle characterization found only subtle differences between the PEV types, which were insufficient to explain their different functional immunomodulatory profiles. Constitutively released PEVs, formed in the absence of an exogenous activator, displayed a disparate activation profile from the receptor induced PEVs.

Our findings underscore that PEVs are tunable through receptor-mediated activation. To truly comprehend their role(s) in mediating platelet functions among immune cells, conducting functional assays is imperative.

**Graphical Abstract:** 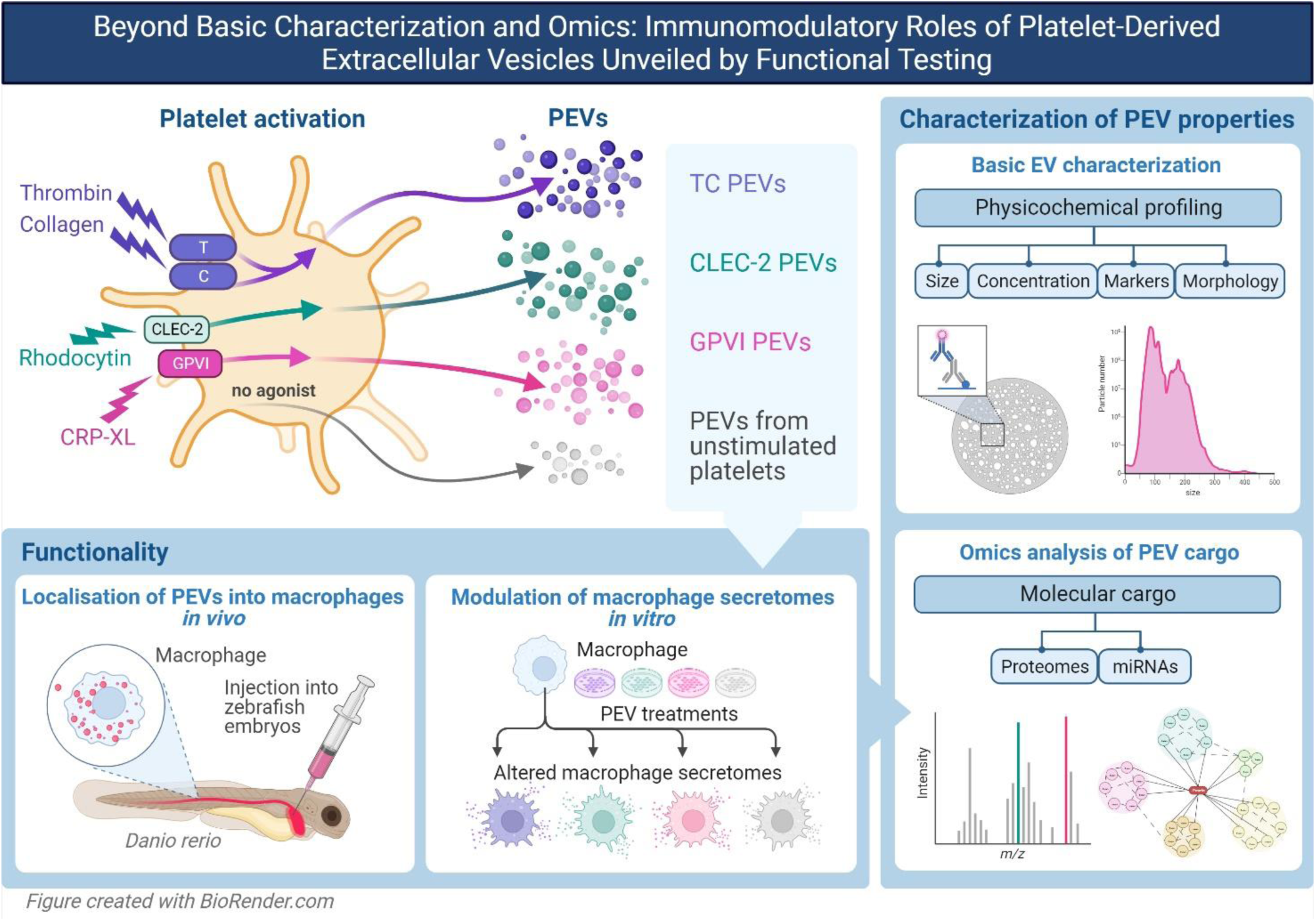

## 1. Introduction

Platelets, anucleate cells originating from megakaryocytes, exhibit heterogeneity in molecular content and functional profiles. They express receptors on their plasma membrane, which enable versatile signaling-dependent activation endpoints ranging from adhesion to thrombus formation and procoagulant transformation^1^. Platelets are also recognized for their roles in immunothrombosis^2^ and immune-mediated inflammatory diseases^3^. Upon activation, platelets release variable classes of soluble bioactive molecules and extracellular vesicles (PEVs)^1^. Previous studies have shown that platelets are probably the most tunable cells regarding different biogenetic pathways for EV formation, and that the molecular cargo of PEVs, such as their proteome^4,5^, miRNAs^6^, and even the content of cytoplasmic organelles like mitochondria^7^, can vary depending on receptor-mediated activation or the cellular environment, e.g. depending on the shear conditions in circulation^8^. Early electron microscopy (EM) analysis of activated platelets described two populations of PEVs: microvesicles and exosomes^9^. Subsequent EM analyses of PEVs from activated platelets have revealed several morphological classes, including spherical EVs, tubular EVs and large platelet fragments in a size range of 50 - 1000 nm^7,10^ illustrating the exceptional capacity of platelets to generate heterogenous EV subpopulations^11–13^. However, the functionality of these different PEVomes (i.e., all platelet EVs) remains poorly understood.

EVs containing the specific platelet and megakaryocyte integrin subunit CD41 (combined with CD61) constitute ~30 - 52% of all circulating EVs depending on the used analytical method ^14,15^. Although some of these EVs may be of megakaryocyte-origin^16^, elevated levels of circulating PEVs have been associated with pathological conditions with immune-mediated inflammatory mechanisms, such as sepsis^17,18^, autoimmune conditions (lupus, rheumatoid arthritis)^19^, cardiovascular diseases^20^ and cancer^21^. Regarding physiological functions, roles of PEVs have been assigned to coagulation and into recruitment and coordinating “homing” of innate immune cells to the site of vascular injury/infection, whereas their role in coordinating adaptive immune cell functions is less explored^22^.

We hypothesized that to be relevant in different physiological and pathological immune related functions, platelets must be able to control their PEV release to match the activating signal, which makes them exceptionally agile cells regarding EV-production, and, in turn, these PEVs an elegant manner to further disseminate immunomodulation. Thus, we explored the possibility to selectively tune the PEV release by targeting two immunoreceptors containing the immunoreceptor tyrosine-based activation motif (ITAM), glycoprotein VI (GPVI) and C-type lectin-like receptor 2 (CLEC-2), which share the same downstream signaling, but are differently engaged *in vivo*^23^. Regarding inflammatory diseases, GPVI is a master regulator of PEV formation e.g. in rheumatoid arthritis *in vivo* in mice^19^ and is currently a novel drug target for acute ischemic stroke and atherosclerosis^24^. GPVI is activated by collagen and fibrin/fibrinogen^25^ in a thrombus. In contrast, CLEC-2 is a receptor for viruses such as Dengue^26^ and human immunodeficiency virus-1^27^, and can also be activated by podoplanin^28^, which is relevant for cancers, such as melanoma^29^. Importantly, these two ITAM-receptors share signaling pathways which are also common for key immunoreceptors in many immune cells including myeloid, plasma dendritic, B- and T-cells^30,31^. Thus, findings of shared or different properties of PEVs formed upon the engagement of GPVI and CLEC-2 could be of general interest for immune-related inflammatory functions. For this study, we employed the following specific receptor agonists: GPVI PEVs were induced by collagen related peptide (CRP-XL)^32^ and CLEC-2 PEVs by snake venom toxin, rhodocytin^33^. As a well-established PEV-inducer, we employed thrombin and collagen co-activation to engage all platelet thrombin and collagen receptors (TC PEVs)^4,34^, which generates a signal relevant to e.g., cardiovascular pathologies and vascular injury. For comparison, we also isolated PEVs from unstimulated platelets (US PEVs).

To assess the immunomodulatory profiles of PEVs, we performed functional assays *in vivo* and *in vitro*. We first show that PEVs interacted with macrophages *in vivo* after an intravenous injection into zebrafish embryos as a model of the vertebrate reticuloendothelial system^35–37^. We then explored the differences between the four PEV types using a human model of macrophages *in vitro*. Next, we addressed the cargo changes in the PEVs by comparing proteins and miRNAs to identify molecular signatures that could explain the functional differences. We also employed the current state-of-the-art basic EV characterization methodologies, including nanoparticle tracking analysis (NTA), microfluidic resistive pulse sensing (MRPS), single-particle interferometric reflectance imaging sensing (SP-IRIS) and transmission electron microscopy (TEM) to profile the PEVs.

## 2. Materials and Methods

### 2.1 Induction and isolation of platelet-derived extracellular vesicles from platelets

A workflow is given in Figure S1. Platelets were isolated from fresh surplus platelet concentrates (compiled from four donors) obtained from Finnish Red Cross Blood Services (Helsinki, Finland) as described earlier^4^. Platelet concentration was measured with the Beckman Coulter T-540 (Beckman Coulter Inc., Brea, CA, USA) hematology analyzer and adjusted to 250 × 10^6^ per mL with Tyrode-HEPES buffer supplemented with 1 mM MgCl_2_, 2 mM CaCl_2_, and 3 mM KCl (physiological cation levels), matching an average physiological platelet concentration in plasma. Platelets were activated by addition of either i) 2 µg/ml collagen (HORM, Takeda Pharmaceuticals Co., Tokyo, Japan) and 0.2 U/ml thrombin (Enzyme Research Laboratories Ltd., Swansea, UK) (TC)^4^, ii) 10 µg/ml CRP-XL (CambColl Laboratories, UK) (GPVI)^32^ or iii) 10 µg/ml rhodocytin (CLEC-2)^33^ for 30 min in +37 °C. Platelets without any added agonist were treated the same to obtain PEVs from unstimulated platelets. To monitor the extent of activation, changes in platelet concentration and platelet surface P-selectin (CD62P) from platelets and EVs and activated CD41/CD61 (PAC1) from platelets were determined with high sensitivity flow cytometry (Figure S2C). Activation time for the generation of PEVs for the comparative studies was optimized by measuring time curves of CD61+ PEVs (15, 30, 60 and 180 min) to ensure sufficient responses for all agonists (Figure S2D). After activation, platelets were sedimented at 2500 × g for 15 min, after which the supernatant was transferred to a fresh tube and centrifuged at 2500 × g for 15 min at RT to ensure removal of platelet remnants^62^. The absence of residual platelets was verified with the hematology analyzer and the supernatants were analyzed directly, or used for EV isolation. For the PEV isolation, supernatant was overlaid onto a 3 ml 60% iodixanol cushion and centrifuged (Thermo Scientific Sorvall WX Ultra) at 100000 g (SW28Ti rotor, Beckman Coulter) for 3 h after which the fraction above the iodixanol cushion was collected and resuspended in 10 ml of DPBS filtered with a 0.1 µm filter unit (Millex VV, Millipore) to remove any residual iodixanol. The PEVs were then concentrated with 10 kDa cut-off ultrafiltration devices (Amicon® Ultra-15 Centrifugal Filter Unit, Millipore) to a volume of 100 µl, and stored at −80 °C.

### 2.2 High sensitivity flow cytometry

Twenty µl of platelet post activation supernatants were incubated with 2.5 µl of phycoerythrin (PE)-conjugated mouse anti-human CD62P (clone AK-2, BD Biosciences), 2.5 µl of PE-conjugated CD61 (clone VI-PL2, BD Biosciences), or 2.5 µl of fluorescein isothiocyanate (FITC)-conjugated anti-human CD41/CD61 (clone PAC-1, BioLegend) for 2 h at room temperature (RT) in the dark. Isotype-matched negative controls (i.e., samples incubated with mouse PE-conjugated IgG1 or FITC-conjugated IgM at the same concentration) and unlabeled EVs were used to exclude non-specific binding of antibodies and background, respectively. After incubation, the samples were diluted 1:10 with 10 mM Hepes buffer containing 140 mM NaCl, pH 7.2 (NH buffer) that was filtered through 0.1 µm filter unit. All samples were analyzed on an Apogee A50-Micro flow cytometer (Apogee Flow Systems Ltd. UK) at a flow rate of 1.5 µl/min. The flow rate was calibrated using Apogee bead mix (Apogee Flow Systems Ltd. UK) by measuring the concentration of 110 nm FITC-labeled PS beads and comparing the measured concentration to that given by the manufacturer. Each sample was measured for 90 s, with triggering at 405 nm (side scatter). The data analyses were performed with FlowJo (v10.7.1; FlowJo). A Rosetta Calibration system (Exometry) was used to calibrate the light scatter signal and to correlate it to particle diameter^38^. Based on the calibration, size gates were set for PEVs (~200 - 1000 nm), cell remnants (1000 - 2000 nm) and platelets (2000 - 3000 nm). Gates for fluorescent signal were set based on isotype controls. Fluorescent events exceeding the gate were determined as positive events. The presented concentrations represent the number of detected events adjusted for the overall sample dilution, flow rate, and measurement duration. The dilution buffer and free antibody in buffer were used as additional controls and subtracted from the results according to MIFlowCyt guidelines^39^.

### 2.3 Nanoparticle tracking analysis (NTA)

Particle concentration and size distribution were measured with NTA from six biological replicates (24 donors) using a Nanosight instrument LM14 (Malvern Instruments Ltd, Malvern, UK) equipped with a violet laser (405 nm, 70mW) and a sCMOS camera (Hamamatsu Photonics, Hamamatsu, Japan). The samples were diluted in 0.1 µm filtered DPBS to obtain 40-100 particles/frame, and five 30 s videos were recorded with camera level 14. The data were analyzed using NTA 3.0 software (Amesbury, UK) with detection threshold at 4 and screen gain at 10. Statistical analysis was performed using Prism 9 software (GraphPad). Statistical significance was determined using Kruskal-Wallis test (p ≤ 0.05).

### 2.4 Microfluidic resistive pulse sensing (MRPS)

Particle concentration and size distribution were measured with MRPS using a Spectradyne nCS1 instrument (Spectradyne, Torrence, CA, USA) equipped with C300 polydimethylsiloxane cartridges to cover a size range of 50 - 300 nm in vesicle diameter. Samples were diluted in filtered 1% Tween-20 in DPBS to obtain 10^7^ - 10^10^ particles/mL and measured immediately with a sample volume of 4 µl. Data was collected for 300 seconds or until a minimum number of 1000 of particle transition events were counted per analysis. Data was pre-processed with the nCS1 Data Analyzer (Spectradyne) by combining events from two technical replicates, by removing electrical noise and buffer background and by applying cartridge-specific filters to exclude false-positive signals based on transit time vs. diameter, peak symmetry, and signal to noise ratio. The results of data-filtering were graphed in 2D scattergrams and inspected for each run. Statistical analysis was performed using Prism 9 software (GraphPad). Statistical significance was determined using Kruskal-Wallis test (*p ≤ 0.05*).

### 2.5 Transmission electron microscopy (TEM)

PEVs were loaded on 200 mesh pioloform carbon coated glow discharged copper grids, fixed with 2% PFA, stained with 2% neutral uranyl acetate, embedded in methyl cellulose uranyl acetate mixture (1.8/0.4%) and viewed with Tecnai 12 (FEI Company, Eindhoven, The Netherlands) at 80 kV.

### 2.6 Scanning electron microscopy (SEM)

Platelet activation was stopped after 20 min by adding 2% glutaraldehyde in 0.1 M Na-cacodylate buffer (NaCaC buffer), pH 7.4. The platelets were then transferred to poly-L-lysine coated coverslips, and fixation was continued for 20 min at RT. After fixation, the coverslips were rinsed twice with NaCaC-buffer for 3 min each, osmicated for 60 min at RT in 1% OsO4 dissolved in 0.1 M NaCaC-buffer, dehydrated in a graded series of ethanol/water (50% - 100%, v/v) and critical point dried. The samples were then platinum-sputtered and examined using a Quanta 250 Field Emission Gun (FEI, Hillsboro, OR) scanning electron microscope at 3 kV.

### 2.7 Single particle interferometric reflectance imaging sensing (SP-IRIS)

PEV samples were analyzed with the ExoView^TM^ Plasma Tetraspanin kit and an ExoView^TM^ R100 (NanoView Biosciences, Boston, MA). The samples were diluted with incubation buffer to a desired concentration (1 - 3 × 10^8^ particles in total) based on NTA measurement. Thirty-five microliters of sample was added directly on the chip and incubated at RT for 16 h. The samples were then stained with a cocktail of fluorescently labelled antibodies containing anti-human CD81 (JS-81, CF555), anti-human CD63 (H5C6, CF647) and anti-human CD9 (HI9a, CF488), washed, dried, and scanned in the Exoview scanner for interferometric reflectance imaging and fluorescent detection. The data was analyzed using the NanoViewer analysis software (NanoView Biosciences), using fluorescent cutoffs as follows: 250 AU for the CF555 channel, 480 AU for the CF488 channel, and 269 for the CF647 channel. Statistical analysis was performed using Prism 9 software (GraphPad). Statistical significance was determined using two-way ANOVA (p ≤ 0.05).

### 2.8 Proteomics

PEV samples (total of 1 × 10^10^ particles) were first dried in a vacuum concentrator (Heto VR-maxi ST, Heto Lab Equipment), and resuspended in 50 nM ammonium bicarbonate buffer (AMBIC), pH 7.8, supplemented with 0.1% RapiGest SF (Waters Inc.). Protein concentrations of the samples were determined with a standard BCA protein assay (Thermo Fisher Scientific, Waltham, USA) and 4 µg of each sample was pipetted into a final volume of 50 µl in AMBIC for the tryptic peptide preparation. Tryptic peptides were prepared using an In-Solution Tryptic Digestion method, according to the following protocol: 5 min denaturation at 95 °C, 30 min disulfide reduction with 5mM Tris (2-carboxyethyl)phosphine hydrochloride (TCEP) solution at 70 °C, followed by 30 min alkylation at RT in the dark with 5 mM iodoacetamide (IAA). Sequencing grade trypsin (Promega) was added 1:50 enzyme:protein for 2 h at 37 °C, and again at the same concentration for overnight incubation at 37°C. After overnight digestion, formic acid was added to a final concentration 0.1%, followed by incubation at 37 °C for 45 min to remove RapiGest SF, after which the samples were centrifugated at 13 000 rpm for 15 min to remove residual debris and transferred into 250 µl autosampler microvials (Thermo Fisher Scientific). Samples were then loaded into an Easy-nLC 1200 (Thermo Fisher Scientific) coupled to an Orbitrap Fusion mass spectrometer (Thermo Fisher Scientific). Peptides (100 ng) were separated using Acclaim PepMap C18 columns (2 µm, 100 Å, 75 mm, 15 cm; Thermo Fisher Scientific). The peptides were loaded in buffer A (5% acetonitrile and 0.1% formic acid) and eluted with a 1 h linear gradient from 10% to 40% buffer B (80% acetonitrile and 0.1% formic acid). Three biological replicates (each in three technical replicates) were sequentially injected with two 15 min wash runs and a 1 h blank run alternated between distinct “treatments”. Mass spectra were acquired using a cycle time data-dependent method with an automatic switch between full MS and MS/MS (MS2) scans every 3 sec. The Orbitrap analyzer parameters for the full MS scan were resolution of 120 000, mass range of 350 to 1800 m/z, and standard AGC target, whereas those for MS2 spectra acquisition were resolution of 30 000, AGC target of 5 × 10^4^ ions, with an isolation window of 2 m/z and dynamic exclusion of 30 sec. Column chromatographic performance was routinely monitored with intermittent injections of 50 fmol of a commercially available BSA peptide mix (Bruker Corporation, MA, USA), as well as evaluating double-wash runs for carry-over peptides.

Protein identification and quantification were carried out with MaxQuant^40^ software package v. 1.6.17.0, with the UniProtKB human FASTA file containing > 86 000 entries with 245 commonly observed contaminant and all reverse sequences. Technical replicates (n = 3) of each sample were matched between runs to transfer identifications between replicates. All other parameters were used in their default settings. Perseus data analysis software^41^ v.1.6.14.0 were used for differential abundance analysis and hierarchical clustering. Abundance values were log_2_ transformed, protein identifications classified as being only identified by site, and reverse sequences and potential contaminants were filtered out from the data frame. Additionally, identifications with zero intensity values in all biological replicates were excluded from the comparison. Missing intensity values of were imputed from normal distribution, and abundance intensities were median normalized. Statistical analysis was performed using Prism 9 software (GraphPad Software, MA, USA). Statistical significance was determined using the Kruskal-Wallis test (p ≤ 0.05).

### 2.9 Proximity extension assay (PEA)

The inflammation-related protein cargo of PEVs (n = 3; 12 donors) was analyzed with the Proximity Extension Assay (PEA) technology^42^ using Olink® Inflammation panel (Table S1). The protein concentration of PEVs were measured with DC assay (Bio-Rad) according to manufacturer’s instructions using BSA as a standard. Samples were normalized to 90 µg/ml prior the analysis. Data is expressed as normalized protein expression (NPX) values on a log2 scale whereby a higher NPX correlates with higher protein expression. Data normalization (against extension control and inter plate control) was performed to minimize both intra-and inter-assay variation. Proteins containing NPX values > 50% below the assay’s limit of detection (LOD) were excluded from the analysis. Values below LOD were replaced by LOD/2 and linearized expression of the log2 scale was used for statistical analysis. Statistical analysis was performed using Prism 9 software (GraphPad). Statistical significance was determined using Kruskal-Wallis test (p ≤ 0.05).

### 2.10 Small RNA analysis

PEV samples (total of 1 × 10^10^ particles, 100 µl) from four biological replicates (12 donors) were diluted to 500 µl with DPBS, and then pelleted at 110000 g for 90 min +4 °C by Optima MAX-XP ultracentrifuge with TLA-55 rotor, k factor 81.3 (Beckman Coulter). RNA was isolated from the pelleted samples with a miRNeasy micro kit (Qiagen) following the manufacturer’s instructions and stored at −80°C for further analysis. The quantity and quality of RNA in the samples were evaluated with the Qubit RNA HS Assay kit (Thermo Ficher Scientific), and Agilent RNA 6000 Pico Chips (Agilent Technologies).

SmallRNA sequencing libraries were prepared following the protocol included in the kit NEXTflex^TM^ Small RNA-Seq kit v3 (Bio Scientific Co). Briefly, total RNA of each sample (between 200 and 10,000 pg) was incubated for 2 min at 70°C, then 3’ 4N adenylated adapter (adapter dilution 1/4) and ligase enzyme were added, and ligation was conducted by incubation of this mix at overnight at 20°C. After excess 3’ adapter removal, 5’-adapter was added alongside with ligase enzyme and the mix was incubated at 20°C for 1 hour. The ligation product was used for the reverse transcription with the M-MuLV Reverse Transcriptase in a thermocycler for 30 min at 42°C and 10 min 90°C. Next, enrichment of the cDNA was performed using PCR cycling: 2 min at 95 °C; 22-25 cycles of 20 sec at 95 °C, 30 sec at 60 °C and 15 sec at 72 °C; a final elongation of 2 min at 72 °C and pause at 4 °C. PCR products were resolved on 8% Novex TBE PAGE gels (Cat. # EC6215BOX, Thermo Fisher Scientific), and one band between 150 bp and 300 bp was cut from the gel. Small RNAs were extracted from the polyacrylamide gel using an adapted protocol, in which DNA was eluted from the gel slices in nuclease free water overnight at RT. Afterwards, libraries were quantified using a Qubit dsDNA HS DNA Kit (Thermo Fischer Scientific) and visualized on an Agilent 2100 Bioanalyzer using an Agilent High Sensitivity DNA Kit (Agilent Technologies). Libraries were sequenced in a HiSeq2500 (Illumina Inc.) by at least 12 million 51-nucleotide reads. Once raw counts were extracted for all the samples, lowly expressed genes were filtered out by running a proportion test, as implemented in the NOISeq package^43^. Normalization and differential expression analysis were performed by using the DESeq2 algorithm^44^. To correct the p-values for multiple testing, the False Discovery Rate (FDR) method was applied. Genes were considered differentially expressed if adjusted p-value < 0.05. miRNAs target prediction was performed through the use of the multiMiR Bioconductor package^45^. In particular, we focused our prediction on databases of validated targets, including mirecords^46^, mirtarbase^47^, and tarbase^48^. The functional profiling of the miRNA target genes was performed by using the clusterProfiler Bioconductor package^49^.

### 2.11 *In vivo* functional assay

#### 2.11.1 Zebrafish lines and maintenance

All zebrafish (*Danio rerio*) work was performed in accordance with Danish and EU legislation under permit 2022-15-0202-00169 from the Danish Animal Ethics Council. Zebrafish were maintained under standard conditions (temperature 28 °C, pH 7.2 and conductivity 700 µS) in recirculating housing systems on a 14 hours light - 10 hours darkness cycle and fed three times daily. Embryos were generated by natural crosses and maintained in E3 medium (5 mM NaCl, 0.17 mM KCl, 0.33 mM CaCl_2_, 0.33 mM MgSO4, 10^−5^% (w/w) methylene blue, 2 mM HEPES, pH 7.2) at 28 °C. Experiments were performed on < 5 days post fertilization (dpf) embryos from established transgenic lines; *Tg(mpeg1:mCherry)^gl23^* as a reporter line for embryonic macrophages^50^, and *Tg(tnfa:EGFP-F)^ump5^*for transcriptional activation of *tnfa*^51^. All fish lines and wild type (AB) used in this study were originally obtained from European Zebrafish Resource Centre (Germany).

#### 2.11.2 Intravenous microinjections of PEVs

All microinjection experiments were performed as previously described^35^. Briefly, zebrafish embryos (2 dpf) were dechorionated, anesthetized in E3 medium with 0.016% (w/v) buffered tricaine (3-amino benzoic acid ethyl ester; Sigma-Aldrich), and embedded in 0.8% (w/v) low-melting-point agarose (BioReagent; Sigma-Aldrich). They were then IV injected via the common cardinal vein^52^ with 3 nl of PEVs along with loading buffer (sterile-filtered, endotoxin-free phenol red solution in DPBS; BioReagent, Sigma-Aldrich) giving a nominal dosage range of 10^5^ - 10^6^ particles per bolus estimated by NTA. Injected embryos were imaged *in vivo* without further handling.

#### 2.11.3 Image acquisition, processing, and analysis

As described previously in more detail^35,53^, all images were acquired using a Zeiss LSM 780 upright confocal microscope (Carl Zeiss) with excitation lasers at 488 nm (EGFP), 568 nm (mCherry) and 633 nm (PEVs labeled with CellTrace Far-red dye). The objective lens used was W Plan-Apochromat 40x/1.0 DIC M27 (Carl Zeiss). All images at the designated tissue area were acquired at 2-µm intervals ensuring optical section overlap in each fluorescence channel to construct *z*-stack images and presented as the maximum intensity *z*-stack projections using Fiji/ImageJ^54^. Bright-field image is shown as a single optical slice at an arbitrary position. PEV colocalization with macrophages was determined using a 3D mask approach based on thresholding on the fluorescence intensity of the macrophage reporter protein and the “Convert to Mask” tool in Fiji/ImageJ as described previously^35^. Briefly, for quantification of the PEV fluorescence overlapping with macrophages, the binary mask created was applied by inverting LUT to the channel for PEV fluorescence. This procedure was performed for all optical slices in a stacked image to exclude PEV fluorescence colocalized in the *x,y* space but not in the *z* axis. All images were processed as the maximum intensity z-stack projections and analyzed using Fiji/ImageJ. “Sequestered PEVs” were defined by thresholding of the PEV signals typically picking only immobilized clusters associated with cells. For quantification of the relative contribution of macrophages to overall PEV sequestration, the area of macrophage-masked PEV signal (> threshold) was divided by the area of total PEV signal (> threshold). Since this analysis was critically affected in minor cases where macrophages moved in and out of the field of view, we excluded embryos by applying a criterion that the over-time average of macrophage-masked PEV area should be > 25 µm^2^.

### 2.12 *In vitro* functional assay

#### 2.12.1 Cells

THP-1 cells (ATCC TIB-202), a human monocytic cell line, were grown in complete RPMI media supplemented with 10% EV-depleted fetal bovine serum and 1% GlutaMAX™ Supplement (Gibco). EV depletion from fetal bovine serum was performed using polyethylene glycol^55^. THP-1 cells were plated to 24-well plate at a concentration of 2 × 10^5^ viable cells per well and incubated for 48 h in 1 ml/well of media supplemented with 50 nM phorbol 12-myristate 13-acetate (PMA) (Sigma-Aldrich) for differentiation into macrophages. Viability of the cells was analyzed by Tryphan Blue exclusion test using Countess Automated cell Counter (Thermo Fischer Scientific). After 24 h resting period with fresh media, the cells (n = 4) were treated: 1) GPVI PEVs, 2) CLEC-2 PEVs, 3) TC PEVs, or 4) US PEVs for 6 h and 24 h. For each treatment, a dose of 2.5 × 10^9^ PEVs/well (2 × 10^5^ cells) was used (n = 4), as determined previously^56^. As a control, four wells were grown without PEV addition for 6 h and 24 h. After the incubation period of either 6 h or 24 h, the media was collected, centrifuged at 500 g for 5 min at 4 °C to remove cellular debris and stored at −80 °C for further analysis of cytokines and chemokines.

#### 2.12.2 Cytokine analysis with ProcartaPlex immunoassay

Cytokine secretion of PMA treated THP-1 cells was analyzed using Cytokine & Chemokine 34-Plex Human ProcartaPlex™ Panel 1A (Invitrogen) and measured with xMAP Luminex^TM^ system (Bio-Rad) according to the manufacturer’s instructions. The kit included granulocyte-macrophage colony-stimulating factor (GM-CSF), monocyte chemoattractant protein (MCP) 1, macrophage inflammatory protein (MIP) 1α, MIP-1β, RANTES (regulated on activation, normal T cell expressed and secreted), stromal cell-derived factor 1 (SDF-1α), gamma-induced protein 10 (IP-10), eotaxin, IFN-γ, interleukin (IL) IL-1α, IL-1β, IL-1RA, IL-2, IL-4, IL-5, IL-6, IL-7. IL-8, IL-9, IL-10, IL-12p70, IL-13, IL-15, IL-17A, IL-18, IL-21, IL-22, IL-23, IL-27, IL-31, tumor necrosis factor α (TNFα), TNFβ, interferon (IFN) α, and chemokine (C-X-C motif) ligand (CXCL) 1 (GRO-α). The samples were pre-diluted 1:2 with RPMI media prior to analysis. Standard curves for each protein were generated with a 5-parameter logistic (5-PL) algorithm, reporting values for both median fluorescence intensity (MFI) corrected with background, and concentration data. Proteins having concentration values within the detection range with all biological replicates in at least one sample group were included for further analysis. Log-transformed MFI values were used for statistical analysis demonstrating advantages for including low and high abundant analytes^57^. Statistical analysis was performed using Prism 9 software (GraphPad). Statistical significance was determined using Kruskal-Wallis test (p ≤ 0.05).

## 3. Results

### 3.1 PEVs are tuned for their immunomodulatory functions

To study how platelets tune their PEVs for immune functions, we employed receptor-specific agonists. We compared two immunoreceptors, GPVI and CLEC-2, which share similar downstream signaling in platelets that is analogous to immunoreceptors in leukocytes. GPVI receptors were activated with CRP-XL (GPVI PEVs), and CLEC-2 receptors with rhodocytin (CLEC-2 PEVs), respectively. Thrombin and collagen receptors (including GPVI) were collectively activated by a co-stimulus (TC PEVs) as a well-established PEV inducing signal. We also isolated PEVs from unstimulated platelets to which no exogenous activator was added (US PEVs). After removal of residual platelets post-activation, PEVs were isolated using iodixanol cushion ultracentrifugation. The workflow for platelet isolation, activation and PEV isolation is detailed in Figure S1 and comparative data of platelet activation by the agonists are given in Figure S2.

To first study the fate of PEVs from activated platelets in the circulation of a living organism, we imaged live zebrafish embryos (2 dpf) that were injected with CellTrace^TM^ far-red membrane labeled PEVs. In all zebrafish experiments, we used the *Tg(mpeg1:mCherry)^gl23^* line for labeling all embryonic macrophages with a fluorescent reporter^50^. Following preliminary screening tests with all the four PEV types, we decided to focus on the two PEV types, GPVI and TC, which had high enough concentrations for quantitative image analysis. The PEVs (roughly 10^5^ - 10^6^ particles per bolus by NTA) were systemically delivered into the common cardinal vein that returns the blood from periphery into the heart (Figure 1A). Both the GPVI and TC PEVs predominantly accumulated in blood-resident macrophages already at 0.5 hours and this pattern remained for 6 hours (Figure 1B, Movies 1 and 2). The injected PEVs were rarely detectable in the circulation, indicating that they were rapidly and mostly sequestered by macrophages. While there was no change in the number of macrophages observed in-frame over time (Figure 1C), there was a gradual decrease of the total fluorescence signal from the sequestered PEVs (Figure 1D). The image analysis supports that the PEVs were mainly accumulated in macrophages, with a relative contribution of macrophages to the total PEV sequestration increasing from ~67% to ~83% and from ~63% to ~68% for the GPVI and TC PEVs, respectively (Figure 1E). To further study the functional role of PEVs that targeted macrophages, we traced transcriptional activation of tumor necrosis factor alpha gene (*tnfa*) using a double transgenic combination of *Tg(mpeg1:mCherry)^gl23^*and *Tg(tnfa:EGFP-F)^ump5^* lines. Within 3 hours following the IV injection of the TC PEVs, macrophages started to induce *tnfa*, indicating their polarization towards a pro-inflammatory phenotype (Figure S3, Movie 3).

**Figure 1.**
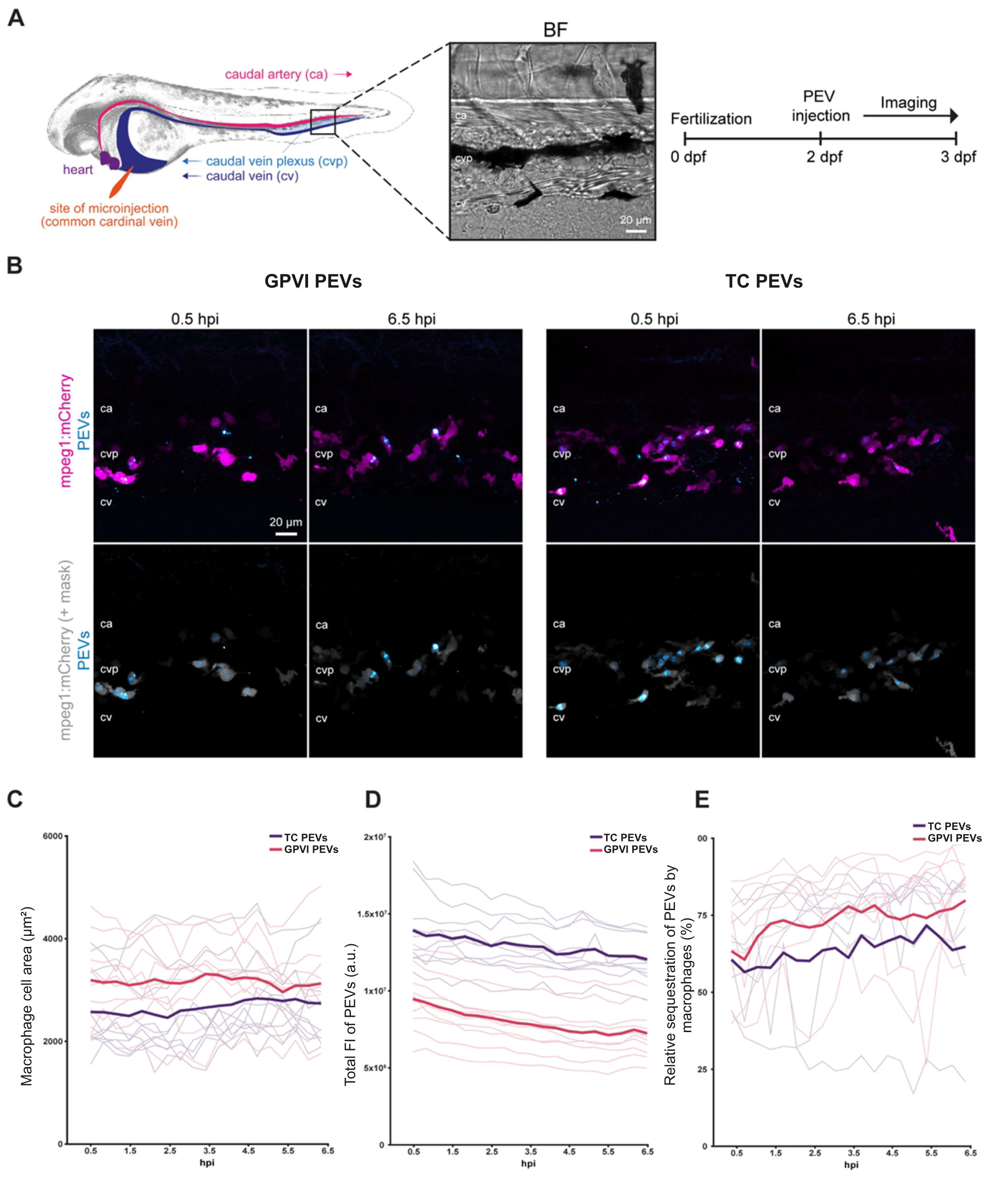
*In vivo* imaging of platelet-derived extracellular vesicle (PEV) sequestration in a zebrafish embryo. **A.** *Tg(mpeg1:mCherry)* embryos at 2 days post fertilization were injected with the fluorescently labeled thrombin and collagen (TC) or GPVI PEVs and imaged from 0.5 h post injection (hpi) to 6.5 hpi. Left panel shows a simplified schematic of the site of microinjection and the main blood vessels including the caudal artery (ca), caudal vein (cv) and CV plexus (cvp) along with a representative bright-field (BF) image of the tissue area imaged. Right panel shows an overview of the experimental setup. **B.** Representative images showing the total PEV signals (cyan, top panel), macrophages (magenta) and PEVs (cyan, bottom panel) with a macrophage-specific mask (gray). Anterior left, dorsal top. **C-E.** Image analysis results. Thin and thick lines represent the individual data and mean values, respectively (*n* = 10 embryos). The relative PEV sequestration by macrophages (E) is the ratio of the macrophage masked PEV area to the total PEV area (*n* = 8 for the GPVI PEVs and *n* = 7 for the TC PEVs after excluding embryos where macrophages were moving in and out of the field of view).

Since *in vivo* imaging indicated that the majority of the PEVs in circulation were taken up by macrophages and could induce *tnfa*, we next wanted to analyze the functional profiles of the differentially induced PEVs more closely on macrophages by comparing their protein secretomes. Macrophages were differentiated from THP-1 cells with PMA and co-cultured with the different PEVs for 6 and 24 hours, after which the macrophage secretomes were analyzed for 34 cytokines relevant to macrophage functions such as polarization (Figure 2A). Macrophages were activated with a concentration of PEVs previously optimized with primary human macrophages^56^. The proteins with concentrations in the detection range of the assay (with all biological replicates in at least one PEV type) were included in the analysis, whereby 16 proteins were analyzed for statistical significance (Table S2). Treatment of macrophages with the different PEV types induced distinct secretomes both at 6 and 24 hours, which were disparate from the control secretomes of the macrophages cultured in the absence of PEVs (Figure 2B). Notably, the secretomes of macrophages treated with the CLEC-2 PEVs were distinguishable already at the 6-hour time point compared to the other PEV types (Figure 2B). While the secretomes induced by the GPVI, TC, and US PEV types differed from the secretome of the untreated macrophages already at 6 hours, distinct profiles among these PEV types emerged at 24 hours, with the separation of the TC PEV-induced secretome from the GPVI and US PEV-induced secretomes. This data indicated that both the kinetics and the profiles induced by the CLEC-2 and GPVI PEVs were different from each other and from the TC PEVs A detailed analysis of the protein changes in the secretomes by comparing the effects of the PEVs from activated platelets (GPVI, CLEC-2 and TC PEVs) with that of the US PEVs showed statistically significant quantitative differences in cytokines. Five out of the 16 cytokines at the 6-hour time point (Figure 2C), and 11 out of the 16 at the 24-hour time point (Figure 2D) were differently secreted after the treatment with the GPVI, CLEC-2 or TC PEVs when compared to the cytokines induced by the US PEVs. At 6 hours, the TC PEVs downregulated five macrophage cytokines (IP-10, IL-10, TNFα, IL-6 and IL-1β), whereas the CLEC-2 PEVs (but only slightly the GPVI PEVs, excluding IL-1β) upregulated these cytokines when compared to the macrophages treated with the US PEVs (Figure 2C). The same trend continued at the 24-hour time point with more cytokines, when the TC PEVs downregulated 10 cytokines (SDF-1α, MIP-1β, MCP-1α, IP-10, eotaxin, IL-1α, IL-10, TNFα, IL-6 and IL-1β) compared to the treatment with the US PEVs. Interestingly, at the 24-hour time point, IL-7, a proliferation cytokine for lymphoid progenitor cells, was upregulated in macrophages by all the PEVs from the GPVI-, CLEC-2- and TC-activated platelets (Figure 2D) when compared to the IL-7 levels induced by the treatment with the US PEVs. To summarize the results, clearly distinct macrophage cytokine profiles from each other were induced by the CLEC-2 and TC PEVs. These profiles were also disparate from the GPVI-induced secretome, which was surprisingly similar to the one induced by the US PEVs.

**Figure 2.**
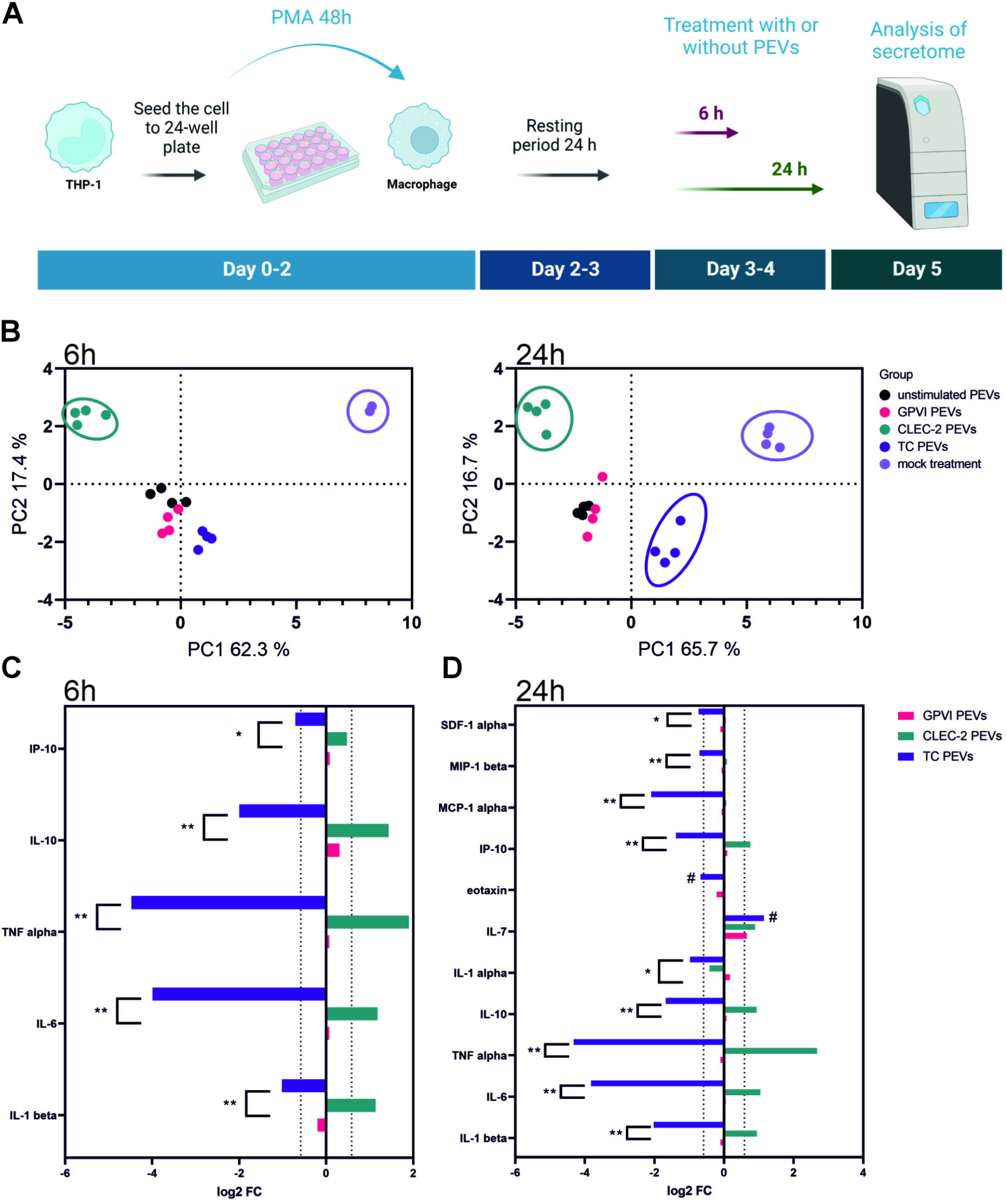
Changes in the macrophage cytokine secretomes induced by the different platelet extracellular vesicle (PEV) types. **A.** Flow chart of the experiment. THP-1 cells were differentiated into macrophages by 48 h incubation with 50 nM PMA. After a resting period of 24 h, macrophages were treated with the different PEV types (n = 4 biological replicates, 12 donors) or left untreated for 6 h or 24 h, after which the media was collected, processed, and analysed for 34 cytokines using a Luminex inflammation panel. Figure created with BioRender.com. **B.** Principal component analysis of the cytokine secretomes of the macrophages at 6 h and 24 h. Secretomes from the mock treated macrophages remained distinct from the PEV-induced secretomes at both time points. At the 6 h timepoint, the CLEC-2 PEV-induced secretome already clustered away from the other PEV-induced secretomes. At the 24 h time point, the TC PEV-induced secretome separated from the other PEV-induced secretomes. Surprisingly, the secretomes from the macrophages treated with the GPVI or US PEVs remained clustered together. **C.** Bar graph showing fold changes (> 1.5 fold change, 0.585 log2 FC) in the fluorescence intensities of cytokines from the GPVI, CLEC-2, and TC PEV-treated macrophages compared to the cytokines from the macrophages treated with the US PEVs at 6 hours. Statistical differences between the GPVI, CLEC-2 or TC PEV-induced secretomes are marked with asterisks (* *p*-value ≤ 0.05, ** *p*-value ≤ 0.01, Friedman test), and the differences between the GPVI, CLEC-2 or TC PEV-induced secretomes compared to the secretome induced by the US PEVs are marked with a pound sign (#). **D.** Bar graph showing fold changes (> 1.5 fold change, 0.585 log2 FC) in the fluorescence intensities of cytokines induced by the GPVI, CLEC-2, or TC PEV-treated macrophages compared to the cytokines from the macrophages treated with the US PEVs at 24 h. Statistical differences between the cytokines induced by the GPVI, CLEC-2 or TC PEVs are marked with asterisks (* *p*-value ≤ 0.05, ** *p*-value ≤ 0.01, Friedman test), and the differences between the GPVI, CLEC-2 or TC PEV-induced secretomes compared to the secretome induced by the US PEVs are marked with a pound sign (#).

### 3.2 Only few differences can be detected between the differentially induced PEVs by basic EV characterization

To identify causes for their differential functionality, the four PEV types were analyzed by methods commonly used to characterize EVs. The isolated PEVs were first imaged with TEM to discern any morphological differences but no notable differences were found between the PEV types (Figure S4A). To study the differences in the PEV generation potency between the agonists, particle concentration, size distribution and surface markers of the PEVs were characterized with three orthogonal single-particle methods (Figure S4B-C). The biggest difference among the different PEV-inducing conditions was in the particle numbers by NTA (Figure S4B). Engagement of GPVI induced the highest particle yield (1.9 × 10^9^) per 2.5 × 10^8^ platelets/ml, followed by TC (1.7 × 10^9^) and CLEC-2 (4.1 × 10^8^) activation, all surpassing the number of particles from unstimulated platelets (2.1 × 10^8^) when measured by NTA.

The same potency order was also detected with MRPS, but with less differences (Figure S4B). Notably, the yields from the GPVI and TC activations were very similar, but despite the similarities of the GPVI and CLEC-2 signaling pathways, CLEC-2 was again a less efficient activator than GPVI. Substantial differences were also evident in the donors’ responsiveness to generate PEVs as judged by the large standard deviations of the particle counts (Figure S4B). Finally, the same order of potency was also seen with anti-human CD61 or anti-human CD62P labeling with direct analysis of the platelet activation samples (without PEV isolation) in high sensitivity flow cytometry (Figure S2D).

Next, we explored whether the activation pathway influenced the size distribution of the PEVs. The PEV samples were analyzed using NTA for particles ranging from 70 - 1000 nm, MRPS for particles within the range of 50 - 250 nm, and SP-IRIS to identify CD41-positive particles in the range of 50 - 200 nm (Figure S4C). Particles were sorted into bins and quantitated. The mean and mode diameters (nm ± SD) are summarized in Figure S4 table. The orthogonal particle analyses revealed that, ultimately, no significant differences could be observed in the size distribution profiles, regardless of the different agonists employed for platelet activation. Only a slightly larger mean particle diameter was detected in the TC-induced samples compared to the others, when analyzed with NTA and SP-IRIS (CD41-captured PEVs), but this observation was not replicated by MRPS.

Next, the subpopulation heterogeneity was inspected by SP-IRIS. Antibodies against three common EV tetraspanins, CD9, CD63 and CD81, were used to capture PEVs in addition to the platelet-specific antibody against CD41. The highest number of PEVs was captured with CD9 (range 1.5 × 10^7^ - 5.8 × 10^7^ particles), and lowest with CD81 (4.6 × 10^5^ - 1.2 × 10^6^), whereas CD63 (range 6.2 × 10^6^ - 2.9 × 10^7^) and CD41 (range 8.1 × 10^6^ - 4.1 × 10^7^) were in between the two. Analysis of the co-localization of tetraspanins CD9, CD63 and CD81 on the CD41-captured PEVs (Figure S5A) showed significant differences between the activation modes regarding the co-localization of CD9 and CD9/CD63 with CD41 (Figure S5B). Compared to the PEVs from the non-stimulated platelets, expression of CD9 in the CD41-positive PEVs was downregulated in the GPVI, CLEC-2 and TC PEVs. In contrast, the GPVI and CLEC-2 PEVs displayed upregulation of CD9/CD63 double positive PEVs compared to the PEVs from unstimulated platelets. The same trend was present in the TC PEVs, but the difference was not statistically significant (2-way ANOVA followed by Tukey’s multiple comparisons test). In all PEV types, the co-expression of CD81 with CD41 was negligible.

To summarize, the basic EV particle characterization only revealed differences in particle yields, where the most notable difference among activations was between GPVI and CLEC-2. GPVI was as potent as TC, whereas CLEC-2 weakly induced PEV formation almost comparable to the particle yields from unstimulated platelets. No differences in size distribution or morphology were observed. However, the tetraspanin co-localization profiles of all the receptor-induced PEV types were the same showing e.g. externalization of CD63, and distinct from the profile of the US PEVs.

### 3.3 Profiling the PEV types through protein and miRNA omics analyses reveals only subtle differences, which are mostly quantitative

Next, we wanted to profile the underlying molecular fingerprints of the PEVs to explain the observed functional differences in the macrophages. To investigate the extent of the platelet’s ability to tune the molecular cargo of PEVs dependent on the receptor activation, we performed three complementary analyses: protein mass spectrometry, PEA of proinflammatory proteins and miRNA sequencing.

Firstly, the protein content of the PEVs was analyzed with mass spectrometry. In total, 250 proteins were identified from the samples (Table S3). Out of 250 proteins, 236 were converted to gene ID for further examination. A Reactome pathway analysis^58^ of all proteins showed that, in addition to the expected platelet-related pathways (e.g. R-HSA-114608, R-HAS-76005, R-HAS-76002), several immune system-related pathways (e.g. complement cascade R-HAS-166658, regulation of complement cascade R-HAS-977606, and innate immune system R-HAS-168249 pathways) were in the top 15 enriched pathways (Figure 3A). Gene ontology analysis revealed that several proteins (YWHAE, VCP, C4BPA, ENO1, PARK7, CORO1A, HSP90B1, TUBA1B, TMSB4X, FLNA, YWHAG, PDIA3, HSPA8, HSP90AA1, ACTN1, MSN, NAP1L1, YWHAZ, PDIA4, PKM, ZYX, MYH9, CALR, P4HB, PFN1, ALDOA, PPIB, PPIA and HSPA1A) were associated with the protein corona and RNA binding (GO:0003723). Comparison of the proteome data to the top 100 protein lists of ExoCarta^59^ and Vesiclepedia^60^ identified 36 proteins shared with either one or both of the databases (Figure 3B). These 36 proteins are known EV membrane proteins (e.g. CD9, CFL1 and FLOT1), or cytosolic proteins (e.g. ALDOA and GAPDH) indicating an enrichment of typical EV proteins in the PEVs^61^. Next, the protein abundances of the GPVI, CLEC-2 and TC PEVs were compared to the protein abundance of the US PEVs. A bar graph (Figure 3C) shows the differentially regulated proteins (Table S4) in the GPVI, CLEC-2, and TC PEVs in relation to the proteins in the US PEVs. In the GPVI PEVs, 54 proteins were upregulated and 101 downregulated when compared to the US PEVs. The corresponding numbers were 44 and 17, and 57 and 58 for the CLEC-2 and TC PEVs, respectively. The number of downregulated and upregulated proteins in the GPVI, CLEC-2 and TC PEVs compared to the US PEVs are illustrated in Venn diagrams (Figure 3D), showing minor differences between the proteomes of the receptor-induced PEV types.

**Figure 3.**
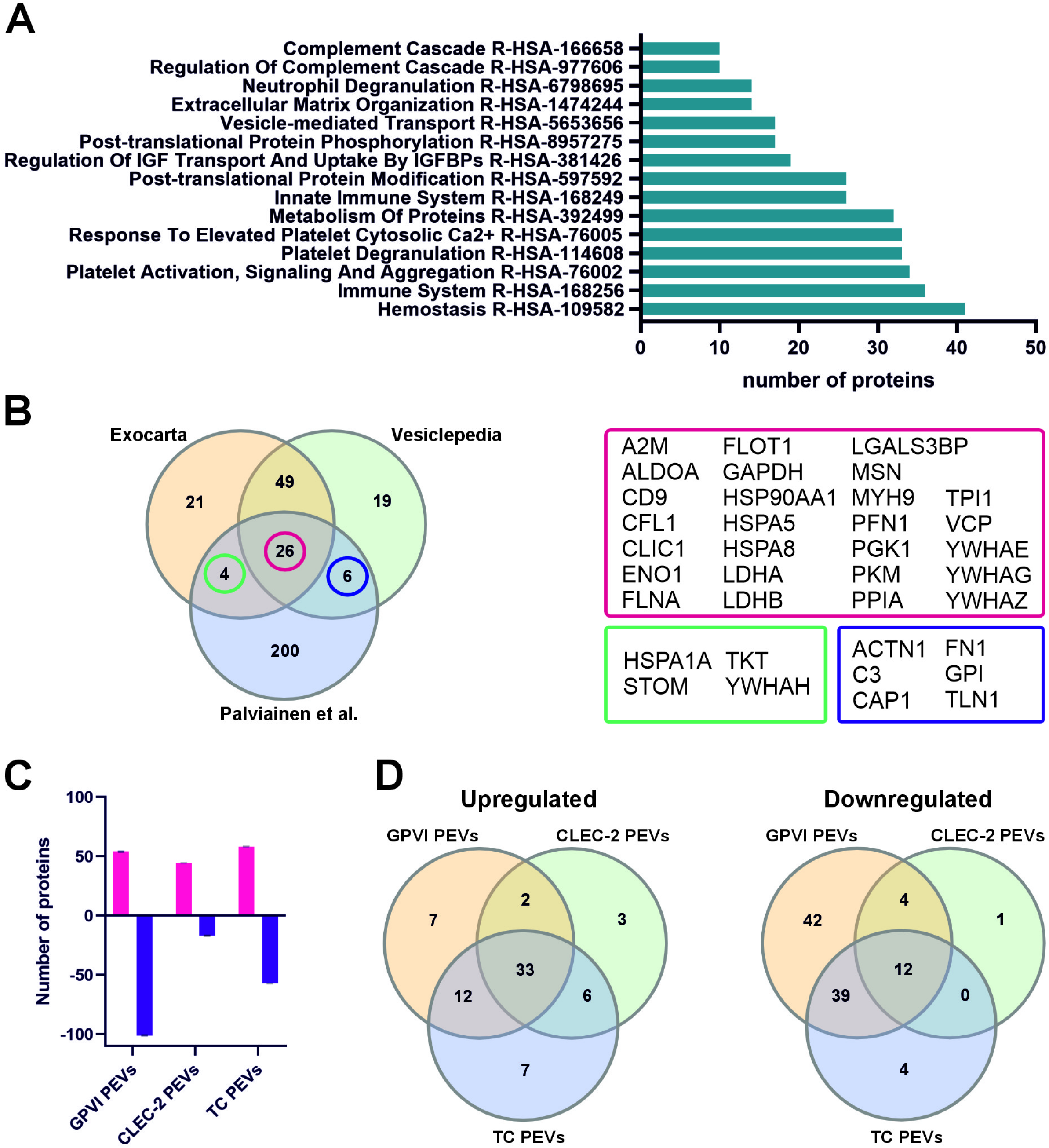
Comparison of the receptor-induced differences in the protein cargo of platelet-derived extracellular vesicles (PEVs) analyzed with mass spectrometry (n = 3) **A.** Bar graph showing the top 15 pathways based on the number of proteins involved in the pathway (≥ 10, adjusted *p*-value ≤ 0.05). **B.** Venn diagram comparing the 240 identified proteins, ExoCarta top 100, and Vesiclepedia top 100 proteins reveals 26 proteins shared with the three datasets. In addition, this dataset shares four proteins with the Exocarta top 100 proteins, and six with the Vesiclepedia top 100 proteins, respectively. **C.** Bar graph illustrating the upregulated (red) and downregulated (purple) proteins when the proteomes of the GPVI, CLEC-2 and TC PEVs were compared to the proteome of the US PEVs. Statistical significance was determined with multiple unpaired t-tests with Benjamini, Krieger and Yekutieli test correction. **D.** Venn diagrams of upregulated and downregulated proteins in the GPVI, CLEC-2 and TC PEVs in comparison to the US PEVs.

A large proportion of the proteins identified in this study have previously been reported as part of the molecular corona of EVs^62,63^, although the PEVs were induced from gel-filtered platelets. To further investigate the corona proteins of the PEVs, the current proteome was compared to the proteomes from two previous PEV proteome studies ^4,63^. A Venn diagram shows the overlap in the identified proteins between the studies (Figure S6), and variations in methodologies are listed in Table S5. Next, gene ontology (GO) enrichment analysis was performed for the shared proteins in all datasets, or between any two datasets. Interestingly, 78 proteins were shared between all three studies despite the methodological differences. The molecular function of GO enrichment analysis revealed that most of the protein functions were related to binding of biomolecules (Figure S6), 48 proteins were shared between our study and Tóth et al.^63^ and 56 proteins between the studies by Aatonen et al.^4^ and Tóth et al. These 104 proteins represent GO classes of functions for biomolecule binding and activity. Finally, shared proteomes between our study and that of Aatonen et al., where platelets had also been activated with an agonist^4^, showed an overlap of 27 proteins with cytokine functions and neurotrophin binding.

The functional assays *in vivo* and *in vitro* implicated the capacity of the PEVs to impact inflammation-related events. Since the proteomic data also showed upregulation of immunity and inflammation-related proteins, we further analyzed PEA inflammation panel proteins in the PEVs. PEA allowed quantitative comparisons of 80 proteins between the four PEV types (Table S1). After exclusion of NPX values 50% below the assay’s limit of detection, 41 proteins remained for quantitative comparison (Table S6). Volcano plots show the differentially expressed inflammation proteins in the GPVI PEVs (Figure 4A), CLEC-2 PEVs (Figure 4B), and TC PEVs (Figure 4C) in relation to the US PEVs. In the GPVI PEVs, nine proteins were upregulated, and two proteins downregulated in comparison to the US PEVs. In the CLEC-2 PEVs, eight proteins were upregulated and 11 downregulated, whereas in the TC PEVs eight proteins were upregulated and two downregulated in comparison to the US PEVs. A Venn diagram of the upregulated proteins (Figure 4D) shows seven proteins shared by at least two PEV types: CCL11, CXCL1, CXCL10, CXCL11, IL8 (CXCL-8), MCP4 (CCL13), and fibroblast growth factor 21 (FGF21) (Figure 4E). A Venn diagram of the downregulated (Figure 4F) proteins reveals that adenosine deaminase (ADA), urokinase-type plasminogen activator (uPA) and CD40 (Figure 4G) were shared between at least two PEV types, whereas up- or downregulated proteins that were present only in the GPVI, CLEC-2 or TC PEVs are shown in Figure S7.

**Figure 4.**
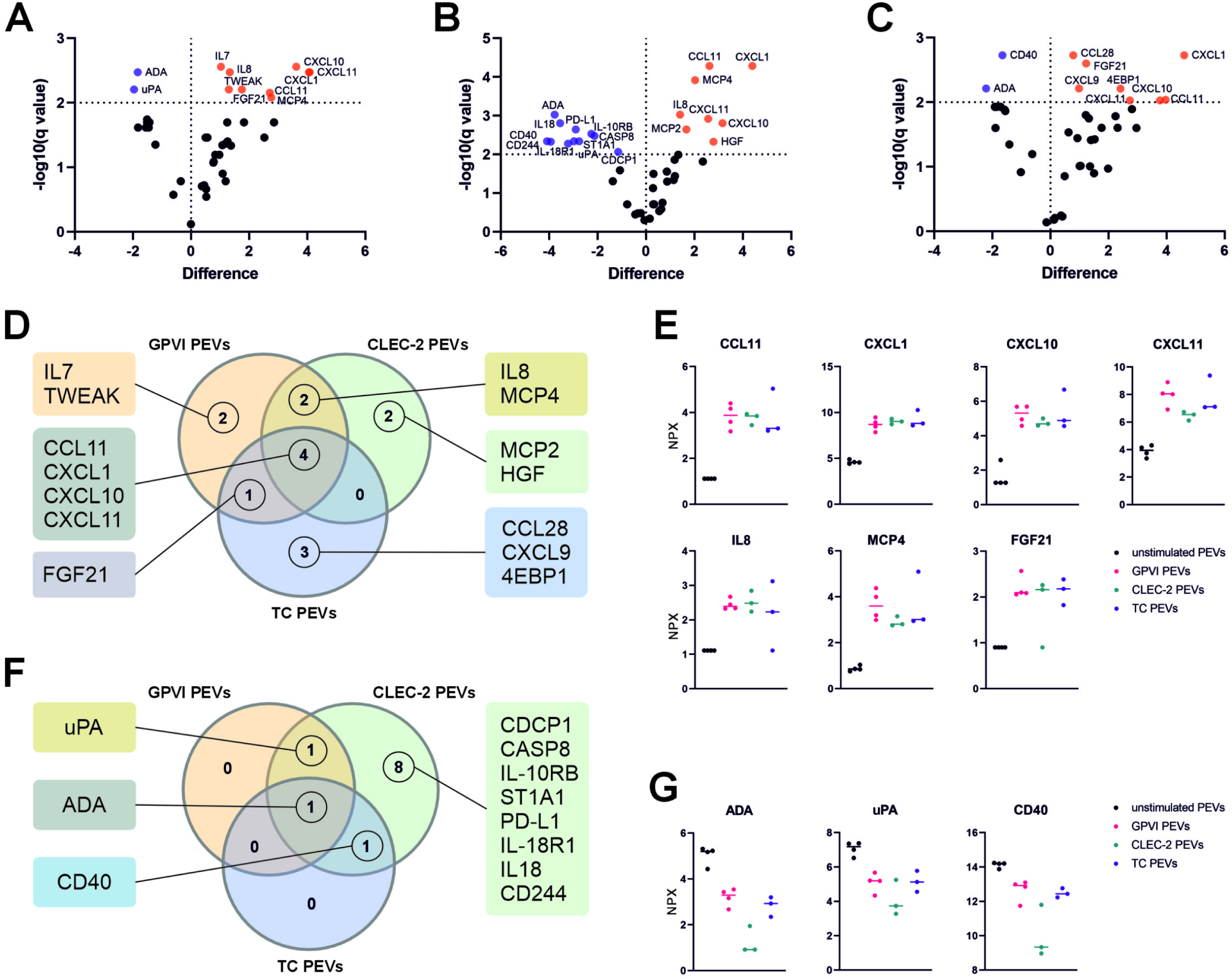
Quantitative analysis of the targets of the inflammation panel by proximity extension assay (PEA) revealed distinctive receptor-dependent tuning of the platelet-derived extracellular vesicle (PEV) proteins. PEVs isolated by iodixanol cushion ultracentrifugation were analyzed for 80 inflammation panel targets. The GPVI (n = 4), CLEC-2 (n = 3) and TC PEVs (n=3) were compared to the PEVs from unstimulated platelets (US PEVs, n = 4). The n values refer to biological replicates, each representing 4 donors. Statistical significance was determined with multiple unpaired t-tests with Benjamini, Krieger and Yekutieli test correction. Results are presented as volcano plots for each PEV type displaying the differently expressed inflammatory proteins compared with those in the US PEVs. Proteins are graphed by difference in means (SO 0.1, x axis) and significance (FDR q < 0.05, y axis). Proteins in orange are upregulated, and in purple downregulated compared to the proteins in the US PEVs. **A.** Volcano plot of the GPVI PEVs compared to the US PEVs shows nine upregulated and two downregulated proteins. **B.** Volcano plot of the CLEC-2 PEVs compared to the US PEVs shows eight upregulated and 11 downregulated proteins. **C.** Volcano plot of the TC PEVs compared to the US PEVs shows eight upregulated and two downregulated proteins. **D.** Venn diagram of significantly upregulated proteins in the GPVI, CLEC-2 and TC PEVs. **E.** Plots illustrating the median expression of seven inflammatory proteins, CCL11, CXCL1. CXCL10, CXCL11, IL8 (CXCL8), MCP4 (CCL13), FGF21, which were upregulated in at least two PEV types. **F.** Venn diagram of significantly downregulated proteins in the GPVI, CLEC-2 and TC PEVs compared to the US PEVs. **G.** Median expression of three inflammatory proteins, ADA, uPA and CD40, which were downregulated at least in two PEV types. P-values were calculated with multiple unpaired t-tests with Benjamini, Krieger and Yekutieli test correction to control the FDR.

Altogether, the comparison between the PEVs from ITAM-receptor activated platelets revealed only quantitative differences, which were mostly downregulated proteins in the CLEC-2 PEVs.

Due to the richness of RNA-binding proteins in the PEVs, we finally analyzed the miRNA content of the PEVs as a possible mechanism of action for their different functionalities. From the four PEV types and platelets, 541 miRNAs were identified by sequencing (Table S7). Although the platelet miRNAs were clearly separated from the PEV miRNAs by principal component analysis (Figure 5A), contrary to our hypothesis, no clear clustering was found among the miRNAs from the US, GPVI, CLEC-2 or TC PEVs. When the differentially expressed genes (DEGs) were analyzed using the DESeq2 algorithm^44^, and the p-values were corrected for multiple testing with False Discovery Rate (FDR) method, 414 DEGs were found to be shared among the PEVs (Figure 5B). Next, the DEGs were compared by their target genes to reveal the impacted pathways. The top ten pathways affected by the shared DEGs from all the PEV types were interleukin-4 and interleukin-13 signaling (R-HSA-6785807), signaling by interleukins (R-HSA-449147), mitotic G1-G1/S phases (R-HSA-453279), cyclin D associated events in G1 (R-HSA-69231), G1 phase (R-HSA-69236), cellular senescence (R-HSA-2559583), transcriptional regulation by Tp53 (R-HSA-3700989), ESR-mediated signaling (R-HSA-8939211), oncogene induced senescence (R-HSA-2559585), and G1/S transition (R-HSA-69206) (Figure 5C). This indicates that the PEV miRNAs may impact many functions including immunomodulatory ones, but no specific profiles could be discerned by the miRNA signatures which could have been assigned with a PEV-generating signal. This conclusion was supported by the final comparison of the miRNAs between the PEVs from the activated platelets (GPVI, CLEC-2 and TC) and the US PEVs. The CLEC-2 PEVs had only three differentially upregulated and one downregulated miRNA when compared to the US PEVs, the GPVI PEVs had 13 up- and nine downregulated, and the TC PEVs 11 up- and eight downregulated miRNAs (Figure 5 D and E).

**Figure 5.**
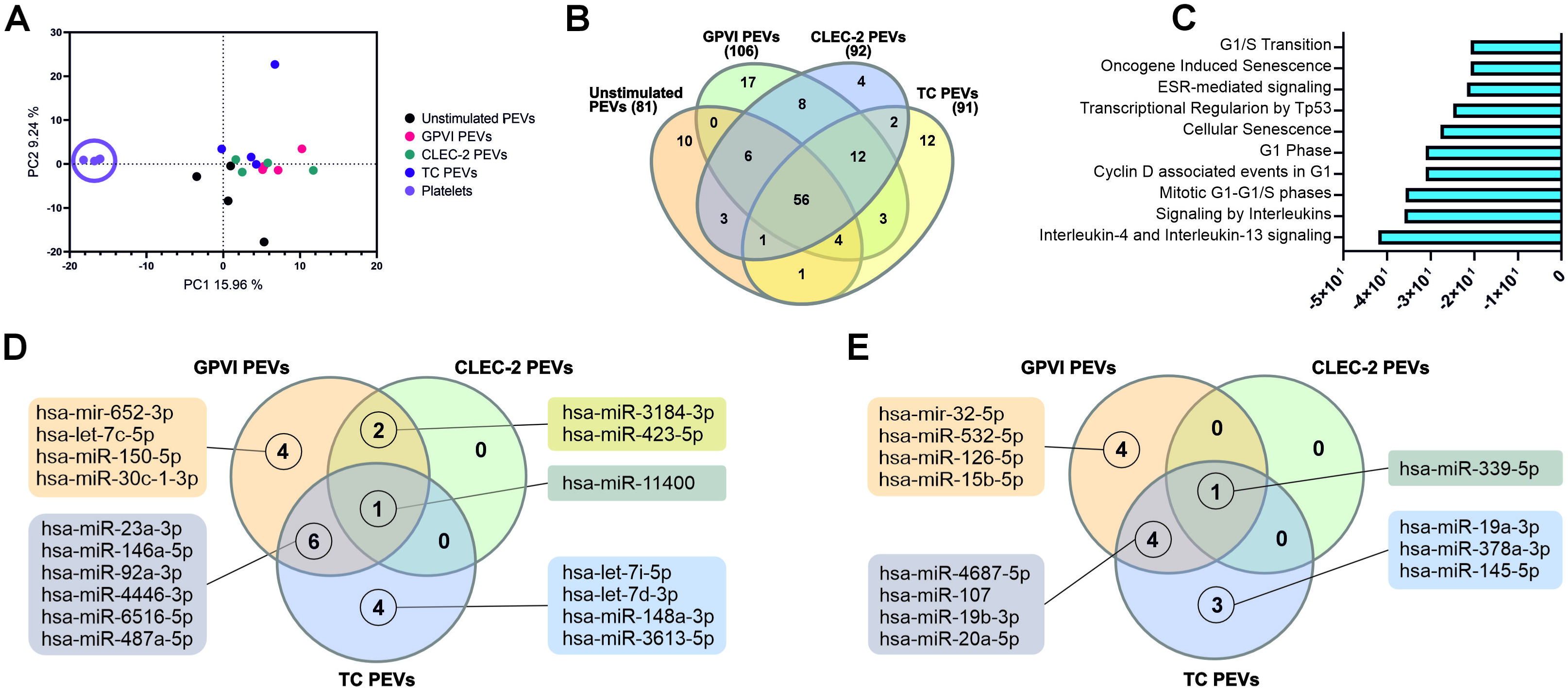
MiRNA content of platelets and the differently induced platelet-derived extracellular vesicles (PEVs) (n = 4; biological replicates representing 16 donors). **A.** Principal component analysis reveals distinct clustering of platelet miRNAs away from the PEV miRNAs, whereas the US, GPVI, CLEC-2 and TC miRNAs overlap with each other. **B.** Venn diagram of the miRNA DEGs in the US, GPVI, CLEC-2 and TC PEVs compared to platelets. **C.** Bar graph of the top ten pathways (p ≤ 0.01) affected by the upregulated miRNAs in the differently induced PEVs compared to platelets. **D.** Venn diagram of the upregulated miRNA DEGs of the GPVI, CLEC-2 and TC PEVs compared to the US PEVs. **E.** Venn diagram of the downregulated miRNA DEGs of the GPVI, CLEC-2 and TC PEVs compared to the US PEVs.

As a summary of the PEV protein and miRNA content analyses, we found that the PEVs were tunable by the platelet activation receptors, and that the US PEVs exhibited their own profile. The differences were, with a very few exceptions, quantitative and not qualitative, which did not allow prediction of a certain functional profile for the different PEVomes.

## 4. Discussion

The *in vivo* imaging of PEVs in zebrafish embryos showed that PEVs were clearly more accumulated by macrophages than by other cells, in particular scavenger endothelial cells (ECs). We find this striking, because a diverse array of nanoparticle types screened by us^35^ and others^36^ were almost always primarily sequestered by scavenger ECs and the contribution of macrophages to the overall nanoparticle clearance was only minor (<20% for 70 nm SiO_2_ nanoparticles)^35^. Indeed, the dominant role of scavenger ECs in nanoparticle sequestration in zebrafish embryos has been reciprocated in rodent models where liver sinusoidal ECs, the functional equivalent in mammals, may contribute more to the hepatic clearance of systemically administered nanoparticles than the Kupffer cells (macrophages)^35–37^. This is also true for some EVs, as zebrafish melanoma EVs (IV injected)^64^ and yolk syncytial layer EVs (secreted *in vivo*)^65^ followed the same homing pattern observed with nanoparticles. This makes our finding on PEVs unique, underscoring that not all EV types behave the same way in circulation. We think that two mechanisms regarding PEVs may play a role here: i) preferential uptake by macrophages previously reported in several *in vitro* studies with human macrophages differentiated from blood monocytes^66–68^, and possibly ii) faster endolysosomal degradation of PEVs following sequestration by the scavenger ECs. The latter scenario is based on the slow disappearance of the total PEV fluorescence (Figure 1D), and similar degradation kinetics have also previously been observed for nanoparticles with a preformed, fluorescently labeled protein corona^53^. Thus, the percentual contribution of PEV uptake by macrophages may be overestimated if scavenger EC sequestered PEVs are rapidly degraded, but even so, our imaging results demonstrate distinct homing of PEVs to macrophages. As there is no counterpart in mature human organs which could be directly represented by embryonic zebrafish macrophages, we rather focused on the primitive and evolutionarily conserved functions of macrophages such as pattern recognition, uptake and inflammation, as demonstrated elsewhere^35–37,64,65^. On the other hand, the cross-reactivity between human PEVs and zebrafish macrophages observed in this study implies conserved recognition mechanisms of PEVs.

The functional *in vitro* macrophage assay showed that PEVs, induced through different platelet receptors, elicit distinct macrophage secretomes. This highlights the significant impact of just subtle molecular tuning within seemingly similar PEVs. Notably, neither basic EV characterization, following established EV field guidelines (such as the MISEV guidelines^61^), nor omics analyses of proteins and miRNA yielded distinct profiles to distinguish the mechanism of action among the different PEVs. In fact, the differences detected between the PEVs generated by the activation of different receptors, such as the two ITAM-receptors, were surprisingly minor. While *in vivo* platelet activation likely involves multiple receptors and signaling pathways, our data underscores two pivotal concepts: i) platelets actively regulate immune cell coordination by tuning the PEVomes, and ii) the analysis of the molecular content of PEVs alone inadequately defines function. These results highlight the multifaceted role of EVs as complex messengers, challenging the notion that analysis confined to proteins, miRNA or basic characteristics is sufficient to reveal such variations. Instead, an integration of biomolecules across diverse classes, also encompassing lipids, sugars, and metabolites, is likely instrumental in orchestrating recipient cell programming, and therefore advocates for a broader perspective in molecular analyses and the need of functional assays.

The original objective of this study was to compare two ITAM receptors in platelets, GPVI and CLEC-2^23^ sharing downstream signaling pathways which are also analogous for the immune receptor signaling in myeloid, plasma dendritic, B- and T-cells. Importantly, the engagement of GPVI in circulation takes place under completely different pathophysiological conditions than that of CLEC-2. According to our hypothesis, we expected distinctly different PEVomes upon stimulation of platelets via GPVI and CLEC-2 if platelets were to engage in immunoregulation via PEVs. Indeed, this was the case based on the marked functional differences in macrophage secretomes and the potency differences between activating receptors, as GPVI induced more PEVs compared to CLEC-2. Unsurprisingly, stimulation through any of the receptors increased the PEV yield compared to unstimulated platelets. In contrast, size distributions of the PEVs were invariant with predominantly smaller PEVs (< 100 nm), as per the power law distribution of EVs^69,70^. However, with the methods used (NTA, MRPS or SP-IRIS), we cannot rule out that differences < 50 nm could be detected with methods such as asymmetric flow field-flow fractionation (AF4)^13^. Surprisingly, the main cargo differences between the GPVI and CLEC-2 PEVs were subtle. There was no statistically significant difference in proteins between these two PEV types and only one miRNA (hsa-miR-140-5p) was significantly upregulated in the GPVI PEVs compared to the CLEC-2 PEVs. However, we cannot rule out the possibility that significant differences could exist in the lipidome, metabolome or glycosignatures of these PEV types, since only proteins and miRNAs were analyzed in this study.

A remarkable finding of this study was the functionality of the PEVs that were formed *in the absence of* an exogenous activator, since also the US PEVs induced a distinct macrophage secretome. The US PEVs exhibited higher levels of 12 proteins (compared to the GPVI, CLEC-2, or TC PEVs), nine of which are known constituents of the biomolecular corona of circulating EVs^62,63^ (A2M, APOA1, APOA4, APOB, APOE, HBB, HPX, ORM1, SERPINA3), and the remaining three are membrane proteins in platelets (CD36, FLOT1, FLOT2). This suggests that the US PEVs originate from the platelet plasma membrane, which is further supported by the loss of 101 proteins in the TC and GPVI-induced PEVs when compared to the US PEVs including immunoglobulins, complement components, and heat shock proteins, which are known constituents of the biomolecular corona and of plasma membrane origin. The origin of the US PEVs is also supported by the tetraspanin co-localization profiles: in comparison to the US PEVs, receptor-mediated activation of platelets decreased the percentage of CD41+/CD9+ PEVs and increased the percentage of triple positive CD41+/CD9+/CD63+ PEVs. Our finding is corroborated by a previous study showing an increase in double positive CD41+/CD9+ EVs in plasma and triple positive CD41+/CD9+/CD63+ EVs in serum^71^ (coagulation activates platelets generating more platelet EVs in serum than in plasma^62,72^). Although we cannot fully exclude inadvertent stimulation by e.g., autocrine ADP release during platelet handling, our data did not show platelet activation and the activation time to induce PEVs was purposedly kept short (30 min) to reduce the possibility of PEVs being generated by aging, apoptosis or necrosis. In support of constitutively released PEVs from unstimulated platelets, previous studies have shown liberation of radioactively labeled platelet membrane in rabbit circulation^73^ and generation of PEVs by platelets over time *in vitro*^4,63,74^ and *ex vivo* in platelet concentrates over the course of several days^75,76^. Therefore, we put forward, that platelets, like other cells, constitutively release small amounts of EVs into the extracellular milieu. Such PEVs could have a homeostatic role in a paracrine manner, and therefore not be readily detected in circulation. Homeostatic function of EVs is a newly emerging concept in the EV field^77^.

Our present data corroborate previous data that activation of platelets via different receptors changes the protein and miRNA content of PEVs^4,5,78–80^. The identification of proteins through non-targeted methods is consistent with our earlier research on platelet EVs^4^, and supports prior studies on the biomolecular corona of EVs^62,63^. Comparison of this data with two prior platelet EV datasets^4,63^ revealed that, interestingly, PEVs exhibit a distinctive set of corona proteins. The majority of these proteins are binding proteins (including RNA binding proteins), suggesting that RNA molecules, as previously proposed, may be transported on the surface of PEVs^81,82^. Furthermore, activation with agonists such as the Ca^2+^ ionophore^4^ and the agonists used in this study resulted in the enrichment of proteins involved in cytokine and chemokine functions, as well as neurotrophin binding, indicating that the receptor-mediated activation is the primary driver of immunologically active PEV formation. This observation was further validated with targeted proteomics (PEA) by measuring a panel of inflammation related proteins and by sequencing of the PEV miRNAs.

Within the last years the knowledge of EV heterogeneity and the existence of various subpopulations with variable biogenetic origins has grown explosively. However, there are still very few systematic comparisons of the effect of different activation modes on the EVome (all EVs) from the same cell type. Platelets represent an excellent cell model for studying the impact of different signaling pathways on the PEVome. Therefore, exploration of the tunability of the PEVome in the context of the immune system is relevant for e.g., the possibility to tune the PEVs towards a defined profile, which would be of interest for the development of drug delivery systems or advanced molecular therapeutic products (AMTP) with innate therapeutic capacity^83^. Also, the high interest in the diagnostic biomarker use of PEVs still demands a better understanding of the variability of the PEVome and its relation to the PEV-inducing signal, whether it is e.g., viral by CLEC-2^84^, thrombotic, mediated by thrombin and collagen receptors^85^, or singularly by GPVI.

Our key findings indicate that PEVs are differentially released *in vitro* both through receptor activation and constitutively in the absence of external stimulation. As anticipated, the activated PEVomes can be tuned; however, understanding their functional role and mechanism of action requires more than just comparative physicochemical characterizations or omics approaches. Instead, we emphasize that sensitive functional assays have the potential to unveil the subtle distinctions among various PEVomes. Altogether, this study accentuates the importance of further and specific functional assays in unraveling the role of PEVs as mediators of platelet functions in the continuum of immune cells.

## Acknowledgements

We thank Laura Bárcena and Dr. Monika González for preparing the smallRNAq libraries and sequencing, and Dr. Felix Royo for his advice in the RNA analysis of EVs. We thank the Affinity Proteomics Unit, ScilifeLab, Uppsala for the PEA analysis. We gratefully acknowledge the fish facility at Aarhus University for zebrafish husbandry and Danna Theresa Vo for her contribution to the original artwork of a zebrafish embryo.

We also thank the HiLIFE core services of the University of Helsinki: EV Core FIMM Technology Centre for performing TEM, Biocomplex unit for ultracentrifugation, and the Electron Microscopy Unit of the Institute of Biotechnology supported by Biocenter Finland for providing the EM facilities.

## Data Availability

The mass spectrometry proteomics data have been deposited to the ProteomeXchange Consortium via the PRIDE partner repository with the dataset identifier PXD047323. The RNA-Seq data have been deposited to the GEO NCBI repository with the accession number GSE184577.

## Supplementary information

### Supplementary Movie Legends

Movie 1. Platelet-derived extracellular vesicle (PEV)-macrophage interactions at 0.5 - 6.5 h post injection (hpi). *Tg(mpeg1:mCherry)* embryos were injected with the fluorescently labeled GPVI PEVs and imaged every 20 min starting at 0.5 hpi. Representative movies are shown. PEVs are seen in cyan and *mpeg1:mCherry* macrophages in magenta. Anterior left, dorsal top.

Movie 2. Platelet-derived extracellular vesicle (PEV)-macrophage interactions at 0.5-6.5 h post injection (hpi). *Tg(mpeg1:mCherry)* embryos were injected with the fluorescently labeled TC PEVs and imaged every 20 min starting at 0.5 hpi. Representative movies are shown. PEVs are seen in cyan and *mpeg1:mCherry* macrophages in magenta. Anterior left, dorsal top.

Movie 3. *tnfa*-induction in macrophages by platelet-derived extracellular vesicles (PEVs) at 1 - 14 h post injection (hpi). *Tg(mpeg1:mCherry);Tg(tnfa:EGFP-F)* embryos were injected with the fluorescently labeled TC PEVs and imaged every 30 min starting at 1 hpi. The movie shows sequestration of PEVs and *tnfa*-induction over time. TC PEVs are seen in cyan, *mpeg1:mCherry* in magenta and *tnfa:EGPF-F* in yellow. Anterior left, dorsal top.

## Supplementary Tables

Table S1. List of proteins targeted in the proximity extension assay (inflammation panel).

Table S2. Macrophage secretomes (cytokines and chemokines) after 6 and 24 h of PEV treatment analyzed by a multiplex immunoassay.

Table S3. Table of 250 identified proteins from proteomics analysis of PEVs.

Table S4. Comparison of identified proteins from proteomics analysis of the GPVI, CLEC-2 and TC PEVs.

Table S5. Comparison of platelet and platelet-derived extracellular vesicle methods used in the studies by Aatonen et al., Tóth et al., and the present study.

Table S6. List of proteins identified from the different platelet-derived extracellular vesicles (PEVs) by the proximity extension assay (inflammation panel). Proteins with NPX values < 50% above the limit of detection were included in the analysis.

Table S7. List of the identified platelet and platelet-derived extracellular vesicle miRNAs.

