## Supplementary material for "Beyond Basic Characterization and Omics: Immunomodulatory Roles of Platelet-Derived Extracellular Vesicles Unveiled by Functional Testing": Table S5

**Table S5.** Comparison of methods for platelet isolation, platelet activation and PEV isolation between Aatonen et al., Tóth et al. and Palviainen et al.

|  | Aatonen et al. 2014^1^ | Tóth et al. 2021^2^ | Palviainen et al. |
| --- | --- | --- | --- |
| Platelet source | Fresh platelet rich plasma from donors | Expired platelet concentrate | Fresh platelet concentrate |
| Platelet isolation | Iodixanol gradient -method^1^  Ca^2+^ -free Tyrode-Hepes buffer (137 mM NaCl, 0.3 mM NaH2PO4, 3.5 mM Hepes, 5.5 mM [D]-glucose, pH 7.35) | Iodixanol gradient -method^1^  Ca^2+^ - and Mg-free Hanks´Balanced Salt Solution (HBSS) | Gel filtration method^1^  Ca^2+^ -free Tyrode-Hepes buffer (137 mM NaCl, 0.3 mM NaH2PO4, 3.5 mM Hepes, 5.5 mM [D]-glucose, pH 7.35) |
| Platelet activation | platelet concentration 250 x 10^6^ /mL  +37⁰C for 30 minutes  10 µM Ca^2+^ ionophore in Ca^2+^ -free Tyrode-Hepes buffer supplemented with 1 mM MgCl^2^, 2 mM CaCl^2^,3mM KCl^2^ | platelet concentration 300 × 10^9^ /mL  +37⁰C, 5% CO_2_ for 180 min, with agitation  Ca^2+^ - and Mg-free HBSS | platelet concentration 250 x 10^6^ /mL  +37⁰C for 30 minutes  Collagen (2µg/ml) and thrombin (0.2U/ml), or CRP-XL (10 µg/ml), or rhodocytin (10 µg/ml) in Ca^2+^ -free Tyrode-Hepes buffer supplemented with 1 mM MgCl^2^, 2 mM CaCl^2^,3mM KCl^2^ |
| EV isolation | Centrifugation 5000g 5 min, RT  Centrifugation 11000g 1 min, RT  Centrifugation 2500g 15 min, RT  Centrifugation 20000g 40 min, +4⁰C (MP sample)  Centrifugation 100000g 60 min, +14⁰C (EXO sample)  Store -80⁰C | Centrifugation 4750g for 5 min, RT  Centrifugation 11000g 1 min, RT  Centrifugation 2500g 15 min, RT  Filtration through 0.8 µm filter  Centrifugation 20000g 40 min, +14⁰C, two times  Resuspension to 10 mM HEPES containing 0.9% NaCl, pH 7.4  Snap freezing, store -80⁰C | Centrifugation 2500g 15 min, RT, two times  Iodixanol cushion -method^3^ combined with washing and concentrating the PEVs using Amicon Ultra 15 centrifugal units  Store -80⁰C |

(1)	Aatonen, M. T.; Öhman, T.; Nyman, T. A.; Laitinen, S.; Grönholm, M.; Siljander, P. R.-M. Isolation and Characterization of Platelet-Derived Extracellular Vesicles. *Journal of Extracellular Vesicles* **2014**, *3* (1), 24692. https://doi.org/10.3402/jev.v3.24692.

(2)	Tóth, E. Á.; Turiák, L.; Visnovitz, T.; Cserép, C.; Mázló, A.; Sódar, B. W.; Försönits, A. I.; Petővári, G.; Sebestyén, A.; Komlósi, Z.; Drahos, L.; Kittel, Á.; Nagy, G.; Bácsi, A.; Dénes, Á.; Gho, Y. S.; Szabó-Taylor, K. É.; Buzás, E. I. Formation of a Protein Corona on the Surface of Extracellular Vesicles in Blood Plasma. *Journal of Extracellular Vesicles* **2021**, *10* (11), e12140. https://doi.org/10.1002/jev2.12140.

(3)	Duong, P.; Chung, A.; Bouchareychas, L.; Raffai, R. L. Cushioned-Density Gradient Ultracentrifugation (C-DGUC) Improves the Isolation Efficiency of Extracellular Vesicles. *PLOS ONE* **2019**, *14* (4), e0215324. https://doi.org/10.1371/journal.pone.0215324.
