## Supplementary material for "Beyond Basic Characterization and Omics: Immunomodulatory Roles of Platelet-Derived Extracellular Vesicles Unveiled by Functional Testing": Figure S1

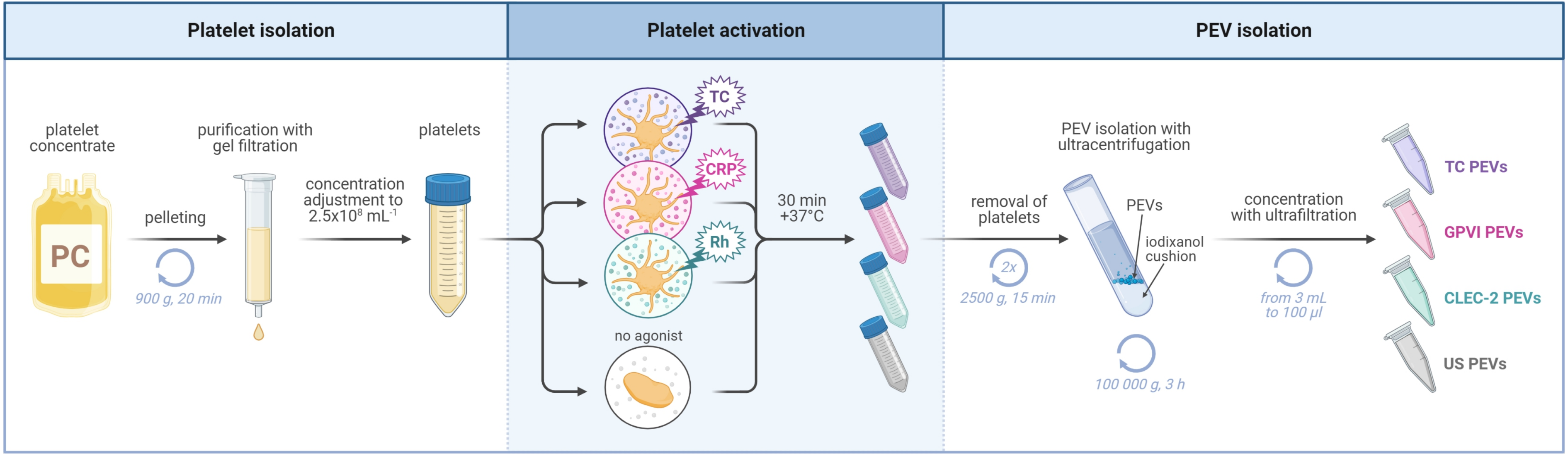


**Figure S1.** Workflow for platelet isolation, activation, and isolation of platelet-derived extracellular vesicles (PEVs). Platelets were isolated from fresh surplus platelet concentrates by gel filtration. Isolated platelets were incubated 30 minutes in +37°C with CRP-XL (CRP), rhodocytin (Rh), and thrombin and collagen co-stimulus (TC), or without any exogenous agonist to produce GPVI, CLEC-2, TC and unstimulated (US) PEVs, respectively. After careful platelet removal, PEVs were isolated with iodixanol cushion ultracentrifugation, washed, resuspended and concentrated with ultrafiltration for downstream analyses. Figure created with BioRender.com.
