## Supplementary material for "Beyond Basic Characterization and Omics: Immunomodulatory Roles of Platelet-Derived Extracellular Vesicles Unveiled by Functional Testing": Figure S2

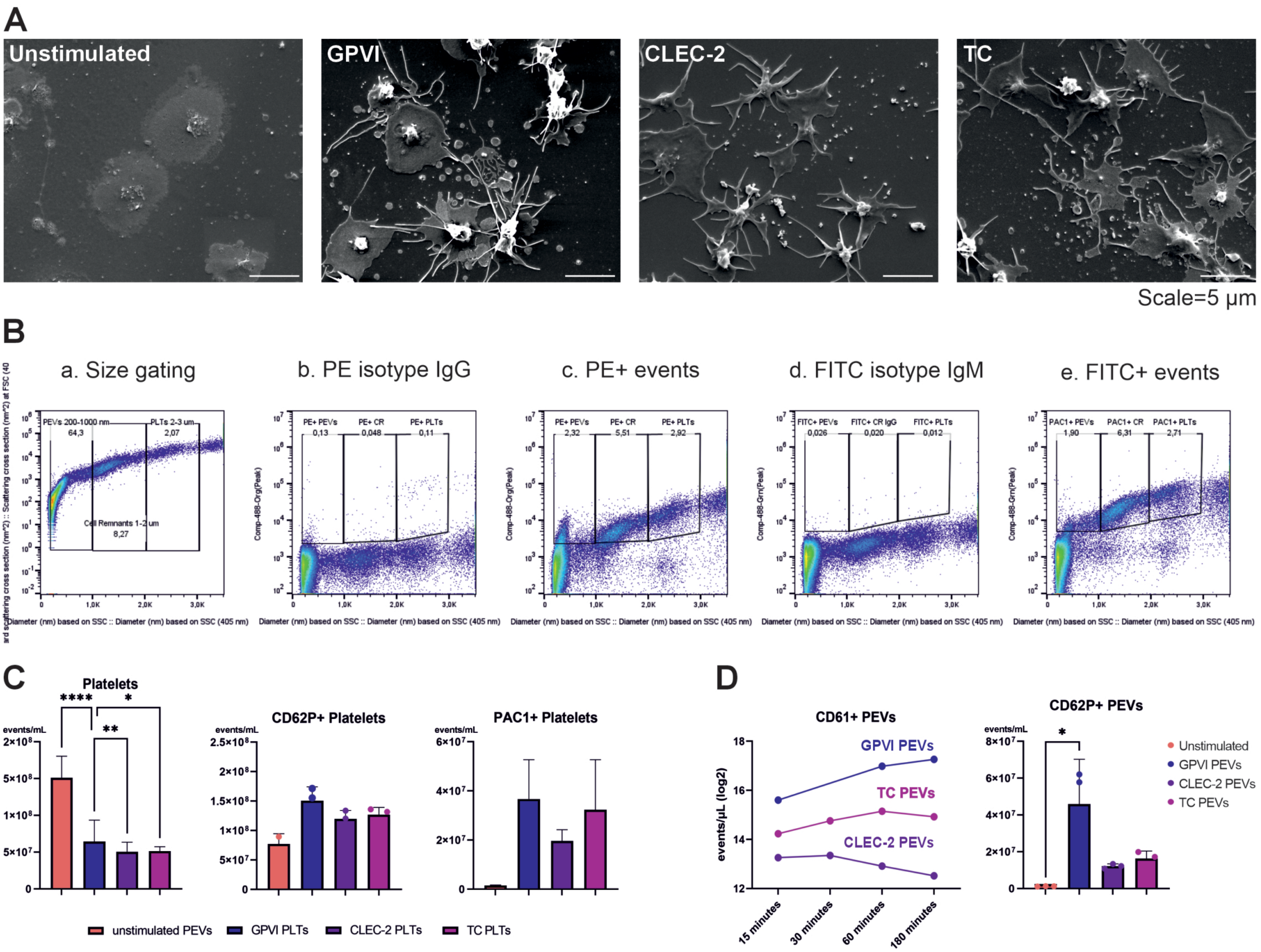


**Figure S2.** Visualization of platelets and platelet-derived extracellular vesicles (PEVs) with scanning electron microscopy (SEM) and high sensitivity flow cytometry (FC). **A.** SEM electron micrographs of unstimulated platelets and platelets activated via GPVI, CLEC-2 and thrombin and collagen co-stimulation (TC). Micrographs show typical morphology changes of activated platelets and PEV shedding compared to the unstimulated platelets. **B.** Gating strategy for PEVs, cell remnants (CRs), and platelets (PLTs) in FC analysis. Representative scatter plots are shown for a. unstained, b. mouse IgG1-PE, c. anti-human CD62P, d. mouse-IgM-FITC, e. mouse anti-human PAC1 stained PEVs, CRs and PLTs. PEVs, CRs and PLTs were detected based on particle diameter derived from light scatter data. Size calibration was done with the Rosetta calibration system (Exometry, The Netherlands) with a PEV diameter size gate of 200-1000 nm, CR diameter gate of 1000-2000 nm and PLT gate of 2000-3000 nm. Fluorescence gates for PE positivity (PE+) were set based on isotype controls to differentiate between specific and nonspecific binding of antibodies. Positive events (+) were defined as events with fluorescent signal exceeding the threshold. **C.** Platelet activation was monitored with FC by deriving platelet concentration from size-calibrated light scatter data (2000-3000 nm), and by measuring the amount of CD62P+ and PAC+ platelets. Upon receptor-mediated activation, the total number of platelets decreased significantly while the number of platelets positive for different activation markers increased compared to the unstimulated platelets (n = 4). **D.** Activation time for GPVI, TC and CLEC-2 stimulation to induce PEV formation was determined by following the generation of CD61+ PEVs at 15, 30, 60 and 180 - minute intervals by FC. The highest yield of PEVs via CLEC-2 stimulation is observed at 30 minutes, where PEV generation is also significantly increased via GPVI and TC compared to unstimulated control (n = 1). The number of CD62P+ PEVs also increased after the 30-minute treatment with agonists compared to PEVs from unstimulated platelets.
