## Supplementary material for "Beyond Basic Characterization and Omics: Immunomodulatory Roles of Platelet-Derived Extracellular Vesicles Unveiled by Functional Testing": Figure S3

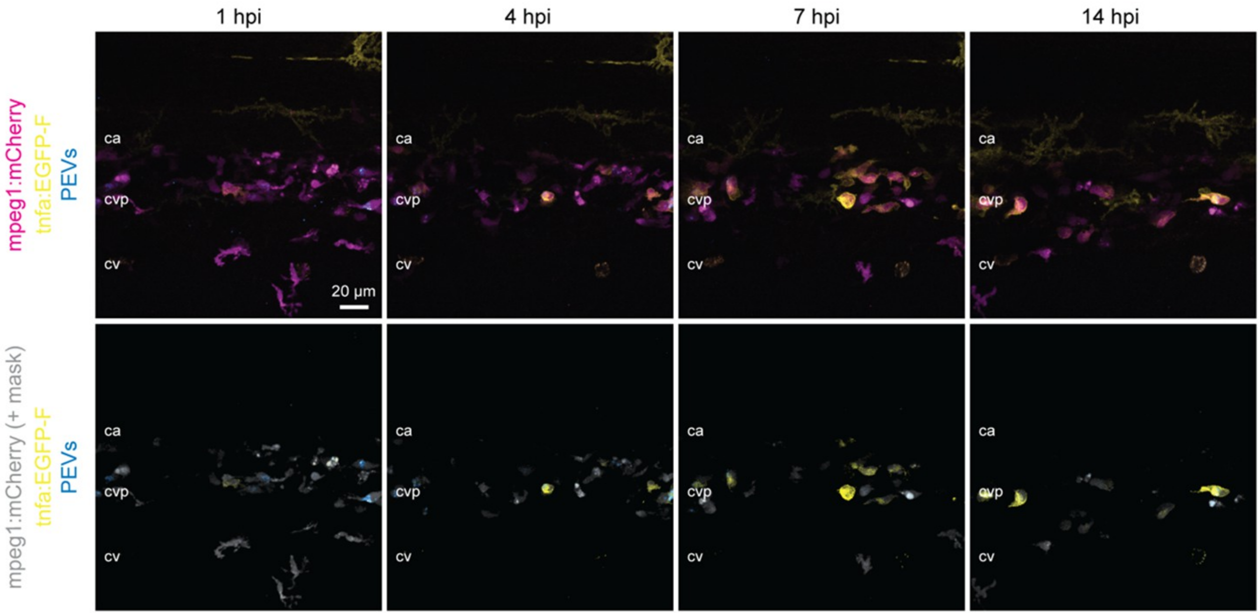


**Figure S3.** Thrombin and collagen (TC) PEVs induce *tnfa* gene expression in macrophages. Two days after fertilization, *Tg(mpeg1:mCherry);Tg(tnfa:EGFP-F)* embryos were injected with the fluorescently labeled TC PEVs and imaged 1 - 14 h post injection. Images showing the time-course (1, 4, 7, and 14 h post injection) of PEV sequestration (cyan) and *tnfa* induction (yellow) with or without a macrophage-specific mask (gray). Anterior left, dorsal top, ca= caudal artery, cvp= caudal vein plexus, cv= caudal vein.
