## Supplementary material for "Beyond Basic Characterization and Omics: Immunomodulatory Roles of Platelet-Derived Extracellular Vesicles Unveiled by Functional Testing": Figure S4

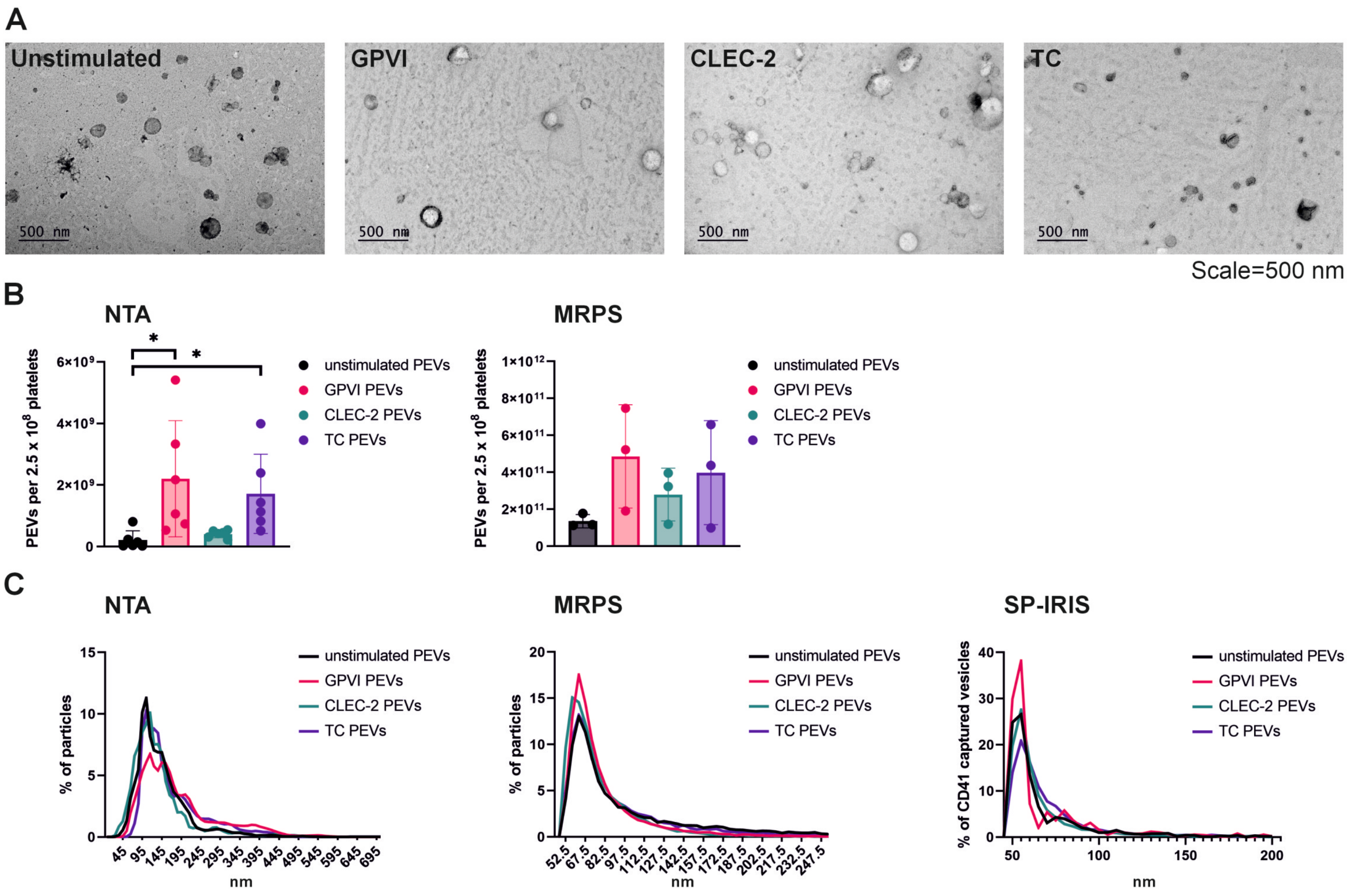


| | US PEVs | | GPVI PEVs | | CLEC-2 PEVs | | TC PEVs | |
|---|---|---|---|---|---|---|---|---|
| | Mean | Mode | Mean | Mode | Mean | Mode | Mean | Mode |
| NTA (n=6) | 140.6 (±17.7) | 106.2 (±5.1) | 152.5 (±24.1) | 123.1 (±26.1) | 146.7 (±13.4) | 112.5 (±18.2) | 170.1 (±18.3) | 126.1 (±9.4) |
| MRPS (n=3) | 65.8 (±1.4) | 62.5 (±5.0) | 70.8 (±1.4) | 60.8 (±2.9) | 66.7 (±5.2) | 60.8 (±5.8) | 65.8 (±3.8) | 60.8 (±5.8) |
| SP-IRIS (n=3) | 60.4 (±1.8) | 55 (±0) | 61.4 (±6.8) | 55 (±0) | 62.2 (±7.1) | 55 (±0) | 66.3 (±8.1) | 55 (±0) |

**Figure S4.** Potency of platelet activation via different receptors in inducing the formation platelet-derived extracellular vesicles (PEVs) and analysis of size distributions. The morphology of the PEVs was compared with transmission electron microscopy (TEM), and particle concentrations were measured with nanoparticle tracking analysis (NTA) and microfluidic resistive pulse sensing (MRPS). Size distributions were studied with NTA, MRPS and single-particle interferometric reflectance imaging sensing (SP-IRIS). **A.** TEM micrographs of the negatively stained GPVI, CLEC-2, thrombin and collagen (TC) PEVs and the PEVs released from unstimulated platelets display typical sizes and morphology for EVs. No differences were observed. **B.** Particle concentrations of the PEVs from unstimulated platelets, and those from platelets activated through the GPVI receptor (GPVI PEVs), CLEC-2 receptor (CLEC-2 PEVs), or with thrombin and collagen receptors (TC PEVs) were measured with NTA and MRPS. Bar graphs represent an average of all analyzed PEV isolates for NTA (n = 6; biological replicates representing 24 donors) and MPRS (n = 3; biological replicates representing 12 donors). PEV concentration is given as PEVs per $2.5^8$ platelets (average concentration in blood) used for the experiment. NTA showed a statistically significant increase in the formation of TC and GPVI PEVs, but not CLEC-2 PEVs, when compared to PEVs from unstimulated platelets ($p > 0.001$; Kruskal-Wallis test followed by Dunn's multiple comparison test). A similar trend was measured with MRPS, although no statistical significance was observed. **C.** Size distributions of the PEVs were measured with NTA, MRPS, and SP-IRIS. The recorded size distributions with NTA and MRPS showed similar size profiles for all PEVs. Data was sorted into 5 nm bins and analyzed with the Kruskal-Wallis test followed by Dunn's multiple comparison test, but no significant differences were found (data not shown). Interferometry-based sizing (SP-IRIS) and counting of PEVs captured with CD41 antibodies also showed similar size distribution profile for all PEVs (n = 3; biological replicates, representing 12 donors). Binned data was analyzed as described above, and no statistical differences were observed (data not shown). Figure table. The mean and mode size of PEVs (nm ±SD) measured with NTA, MRPS and SP-IRIS.
