## Supplementary material for "Beyond Basic Characterization and Omics: Immunomodulatory Roles of Platelet-Derived Extracellular Vesicles Unveiled by Functional Testing": Figure S5

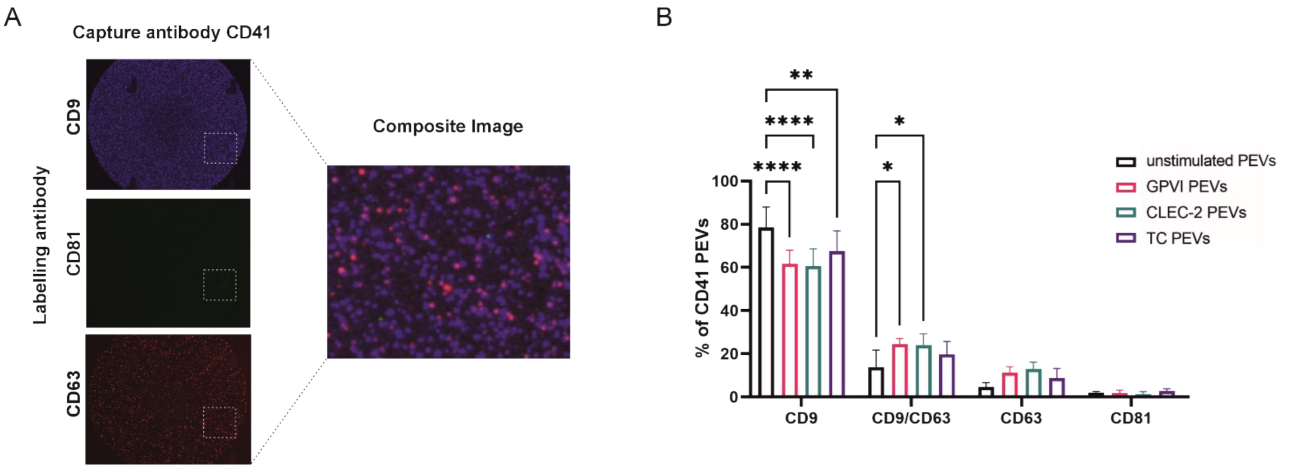


**Figure S5.** Tetraspanin profiles of CD41-captured platelet-derived extracellular vesicles (PEVs) generated via activations of different platelet receptors or in the absence of an exogenous activator. The PEVs were analyzed with single particle interferometric reflectance imaging sensing (SP-IRIS) using the ExoView R100 platform. **A.** Representative image of CD41-captured particles labelled with CD9, CD81 and CD63 antibodies. Source areas for the composite image of co-localized fluorescence markers is shown with a dotted line **B.** Percentages of CD41-positive PEVs labelled with CD9, CD63, and CD81 antibodies. Colocalization profiles of the CD41-captured, CD9-positive and CD41-captured, CD9/CD63-double positive PEVs showed statistically significant differences between the PEV types generated by activating platelets via GPVI (GPVI PEVs), CLEC-2 (CLEC-2 PEVs) and thrombin and collagen receptors (TC PEVs) compared to the PEVs from unstimulated platelets (US PEVs). Graphs represent means ± SD (n = 3; biological replicates representing 12 donors), * $p \leq 0.05$, ** $p < 0.001$, *** $p < 0.0001$ as determined by 2-way ANOVA followed by Tukey's multiple comparisons test.
