## Supplementary material for "Beyond Basic Characterization and Omics: Immunomodulatory Roles of Platelet-Derived Extracellular Vesicles Unveiled by Functional Testing": Figure S6

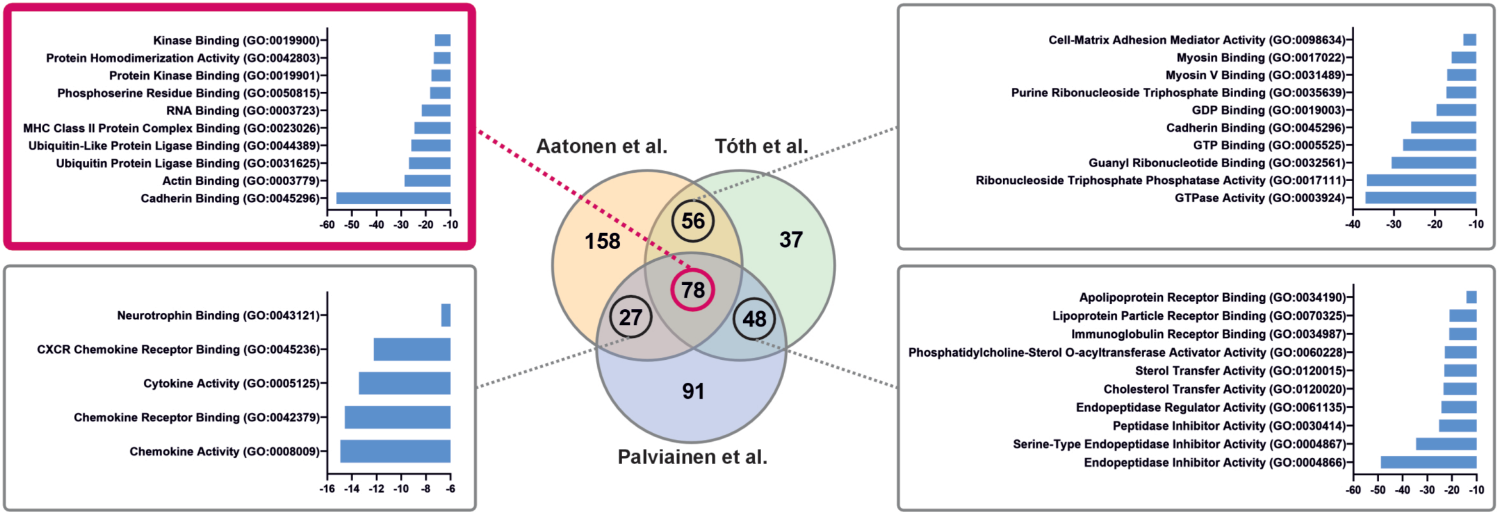


**Figure S6.** Proteomic data comparison of this study with two previous studies of platelet-derived extracellular vesicles (PEVs) by Aatonen et al. and Tóth et al. Venn diagram shows the overlap between the proteomes. Proteins shared by at least two studies were subjected to molecular function of gene ontology (GO) enrichment analysis. Bar graphs represents top 10 molecular functions (p-value ≤ 0.01).
