## Supplementary material for "Beyond Basic Characterization and Omics: Immunomodulatory Roles of Platelet-Derived Extracellular Vesicles Unveiled by Functional Testing": Figure S7

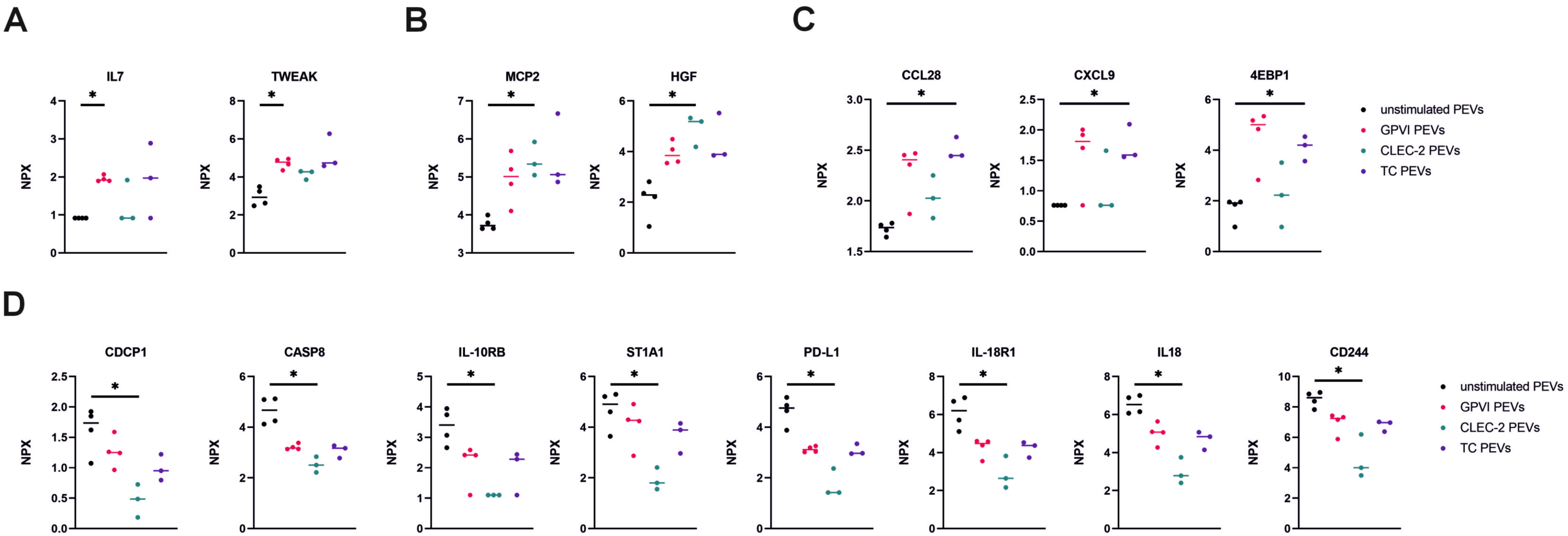


**Figure S7.** Box plots of differently expressed inflammation-related proteins in platelet-derived extracellular vesicles (PEVs) by proximity extension assay (PEA). **A.** Two proteins, IL7 and TWEAK, were significantly upregulated in the GPVI PEVs (n = 4; biological replicates representing 16 donors) compared to the US PEVs (n = 4; biological replicates representing 16 donors). **B.** Two proteins, MCP2 and HGF, were significantly upregulated in the CLEC-2 PEVs (n = 3; biological replicates representing 12 donors) compared to the US PEVs. **C.** Three proteins, CCL28, CXCL9 and 4EPB1, were significantly upregulated in the TC PEVs (n = 3; biological replicates representing 12 donors) compared to the US PEVs. **D.** Eight proteins, CDCP1, CASP8, IL-10RB, ST1A1, PD-L1, IL-18R1, IL18 and CD244 were significantly downregulated in the CLEC-2 PEVs compared to the US PEVs. P-values were calculated with multiple unpaired t-tests with Benjamini, Krieger and Yekutieli test correction to control the FDR, and a p value of ≤ 0.05 was considered significant.
